# Mechanistic Dissection of Conformational Transition of Bicyclic Peptide via Molecular Modeling and Deep Learning

**DOI:** 10.64898/2026.03.12.711417

**Authors:** Ta I Hung, Raghu Venkatesan, Chia-en Chang

## Abstract

Molecular conformations play a critical role in determining molecular properties, such as membrane permeability, binding affinity, and ultimately therapeutic efficacy. Experimental and computational approaches can characterize conformations and provide insight into why certain conformations are thermodynamically preferred over others. However, examining conformations alone may not fully explain why subtle differences, such as a single LEU-to-ILE mutation in a bicyclic peptide, can produce markedly distinct conformational ensembles. Analyzing the transition pathways between conformations further reveals the mechanisms that shape these ensembles. Here, we introduce a deep learning model, termed ICoN-v1, trained in molecular dynamics simulation data to learn the underlying physics that governs cyclic peptide conformational dynamics. We examined hexacyclic peptides with Nuclear magnetic resonance (NMR)-determined structures, and MYC-targeting bicyclic peptides, which are stereo-diversified or have a single LEU-to-ILE mutation. By following the minimum-energy pathway in the latent space constructed by ICoN-v1, conformational transition paths, led by various sets of concerted backbone and sidechain torsional rotations moving in sequence between energy minima, are efficiently generated. Notably, smooth transition pathways that are absent from molecular dynamic output can be observed using ICoN-v1. Our results identify various sets of concerted torsional motions that are nonlinearly combined during conformational transitions and reveal the key residues governing each stage of the transition, thereby elucidating how the observed conformations are generated and informing molecular design.

## Introduction

Understanding how molecules reorganize their conformations is fundamental to decoding how molecules function, especially for cyclic peptides, whose transitions between distinct low-energy states often require highly coordinated, concerted torsional rotations and transient intermediate states ^1,2^. These conformational transitions govern molecular flexibility and enable cyclic peptide to adopt diverse biological properties including membrane permeability, binding affinity, and therapeutic efficacy ^3–6^. Even single-point mutations can substantially alter these pathways, reshape the conformational free-energy landscape and their transition pathway and ultimately affect binding and biological activity. Despite the growing importance of cyclic peptides in drug discovery, with more than 40 cyclic peptide therapeutics approved over the past two decade ^7,8^. The mechanistic details of how cyclic peptides reorganize their torsions during target recognition remain poorly understood. Their structural flexibility enables adoption of multiple out-of-plane conformations that facilitate strong and selective interactions with traditionally “undruggable” protein targets ^9,10^, while cyclization reduces conformational entropy penalties upon binding and partially bound conformations can retain water molecules to reduce desolvation penalties ^3^. In addition, cyclic peptides can enhance membrane permeability by forming intramolecular hydrogen bonds (H-bonds) that shield polar residues ^2,11^. Their large and structured surface areas enable high binding specificity and affinity to their protein targets ^10^. However, capturing these collective dynamics is challenging for both experimental and simulations, limiting rational design of potent cyclic peptides ^12^. A deeper understanding of residue-level motions, concerted torsion angle transitions, and conformational dynamics is crucial to uncover their underlying thermodynamic properties and guide rational design of next-generation therapeutics and uncover their underlying thermodynamic properties.

An experimentally determined structure of a cyclic peptide offers atomic-level structural details but provides limited temporal insights into their motions. For instance, X-ray crystallography typically reveals static conformations. Nuclear magnetic resonance (NMR) spectroscopy offers the ability to observe conformational ensembles in solutions, but it usually provides incomplete represented conformations ^11,13^. To overcome these limitations, computational approaches, particularly molecular dynamics (MD) simulations, have been widely used to explore protein conformations. In addition to classical MD simulations, enhanced sampling techniques sampled molecular conformations more efficiently ^14,15^. However, the resulting trajectories do not inherently reveal and interpret the relationship between peptide sequence and conformational transitions or the underlying physics. Different sets of torsion angles are often required to move in concert to complete a bicyclic ring’s conformational transition processes. However, determining when and how an individual residue contributes to the conformational transition is difficult. Therefore, a more advanced approach is needed to provide mechanistic understanding to efficiently design cyclic peptides.

Artificial intelligence, in particular deep learning (DL), offers an alternative for modeling conformational transitions. Generative models with coarse-grained or protein backbone-only representations utilize a neural network trained on conformations obtained from MD simulations. These models can learn the probability distribution of the training set and allow for sampling novel conformations from the learned distribution, bypassing the sampling bottleneck posed by energy barriers of molecular simulation ^16–19^. However, use of coarse-grained or protein backbone-only representations limits models’ ability to resolve critical side chain, functional-group interactions, and Hbonds networks that determine the conformational senses. Our previous study introduced the model Internal Coordinate Net (ICoN), which leverages vector features to present Bond-Angle-Torsion (BAT) coordinates with a fully connected neural network architecture to achieve efficient conformational sampling for an intrinsically disordered amyloid-β1−42 monomer at full atomistic detail ^20^. Because bicyclic peptides have inherently complex coordinated torsional rotations, existing DL approaches without sophisticatedly incorporating side-chains rotations in the models cannot accurately capture their concerted torsional rotations for dissecting mechanisms of conformational transition of bicyclic peptides.

In this study, we applied a newly developed, physics-based deep learning model, ICoN version 1 (ICoN-v1), to identify transient conformations of cyclic peptides and characterize their conformational transitions pathway. Using classical MD trajectories as the training dataset, ICoN-v1 combines transformer architectures with physics-based loss functions derived from molecular mechanics energy terms to efficiently generate atomistic conformations along smooth and physically interpretable transition pathways between local energy minima in latent space. Analysis of these pathways revealed that intramolecular interactions and coordinated torsional rotations are key drivers of ring puckering and conformational reorganization, providing mechanistic insight into how mutations reshape conformational landscapes and transition mechanisms. Here, we first investigate conformational transition pathways for two hexacyclic peptides, cyclo-(<u>T</u>VGGVG) and cyclo-(<u>V</u>VGGVG), with conformations derived from reported NMR and MD simulations ^21^, to explain why a single-point mutation yields markedly different ring conformations. We then focus on revealing fundamental chemical principles governing conformational transitions for five MYC-targeting bicyclic peptides, reported in our previous publication ^22^ **(Figure 1)**. Although our previous study showed that the different stereo-structures and peptide sequences strongly affect bicyclic peptides conformations and MYC binding, how chirality and individual residues govern distinct conformations remained unknown ^22^. By using ICoN-v1 to generate smooth and physically interpretable transition pathways, we identified energetic contributions of individual residues and the coordinated torsional rotations that drive diverse conformational transitions. This identification explained the effects of chirality and peptide sequences that lead to distinct conformational transitions. The detailed mechanistic insights explain how these molecules move and inform rational design of cyclic peptides.

**Figure 1.**
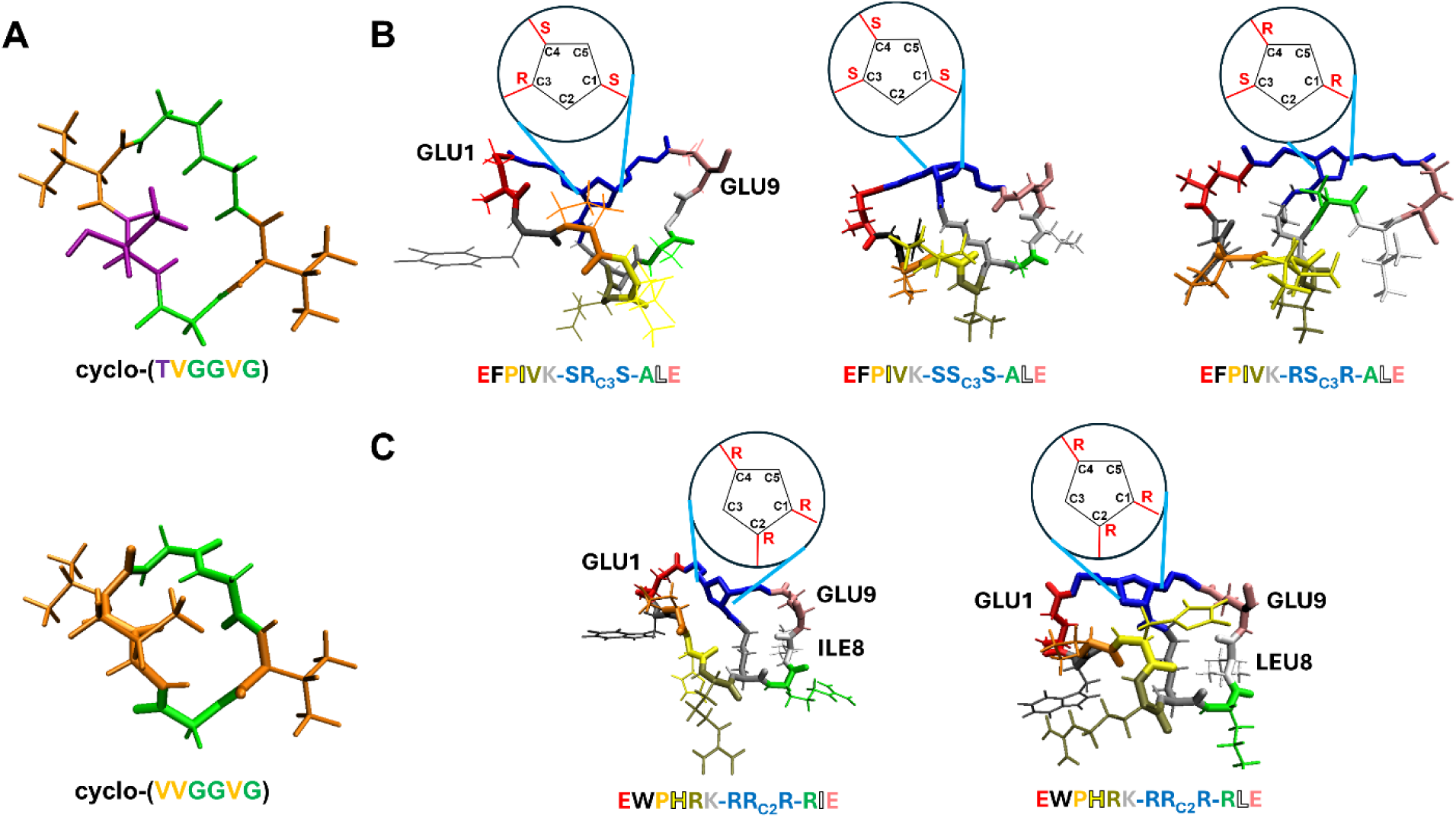
Representative conformation of seven cyclic peptides: **(A)** cyclo-(<u>T</u>VGGVG) and cyclo-(<u>V</u>VGGVG). **(B)** EFPIVK-**SR_C3_S**-ALE, EFPIVK-**SS _C3_S**-ALE, and EFPIVK-**RS_C3_R**-ALE with same amino acid sequence but distinct chirality at C1, C3 and C4 positions. **(C)** EWPHRK-RR_C2_R-R<u>L</u>E and EWPHRK-RR_C2_R-RIE with single point mutation from LEU to ILE at residue 8.

## Methods

### Molecular Structure Representation

#### Bond-Angle-Torsion (BAT) Internal Coordinate

All conformations (peptides and linker) are defined using BAT internal coordinate system. Unlike Cartesian coordinates, BAT coordinates accurately define the circular motion of the dihedral rotation of the molecules ^23^. In fact, BAT is the only coordinate system that can accurately describe the molecular motions when our simulation is performed using molecular mechanics force field (See detail in Supplementary Information for MD simulation protocol).

BAT is defined as sets of root atoms, and the positions of other atoms depend on the previously defined atoms bonded in sequence. For instance, Atom 1 is at the origin; atom 2 is defined by a bond length (b2) and placed on the z-axis; and atom 3 is defined by a bond length (b3), bond angle (a3) and placed on the x–z plane, and thus we eliminate 6 degrees of freedom. The total degrees of freedom of a molecule with N atoms, represented with internal coordinates, has is 3N-6. Atoms 1–3 are termed root atoms for this molecule. Atoms i > 3 are defined by (b_i_, a_i_, □_i_), where t are torsion angles. For example, atom 4 is presented by (b_4_, a_4_, □_4_) (**Figure 2B**). The internal coordinate is then constructed in a tree. Starting from root atoms, all subsequent atoms are defined sequentially.

**Figure 2.**
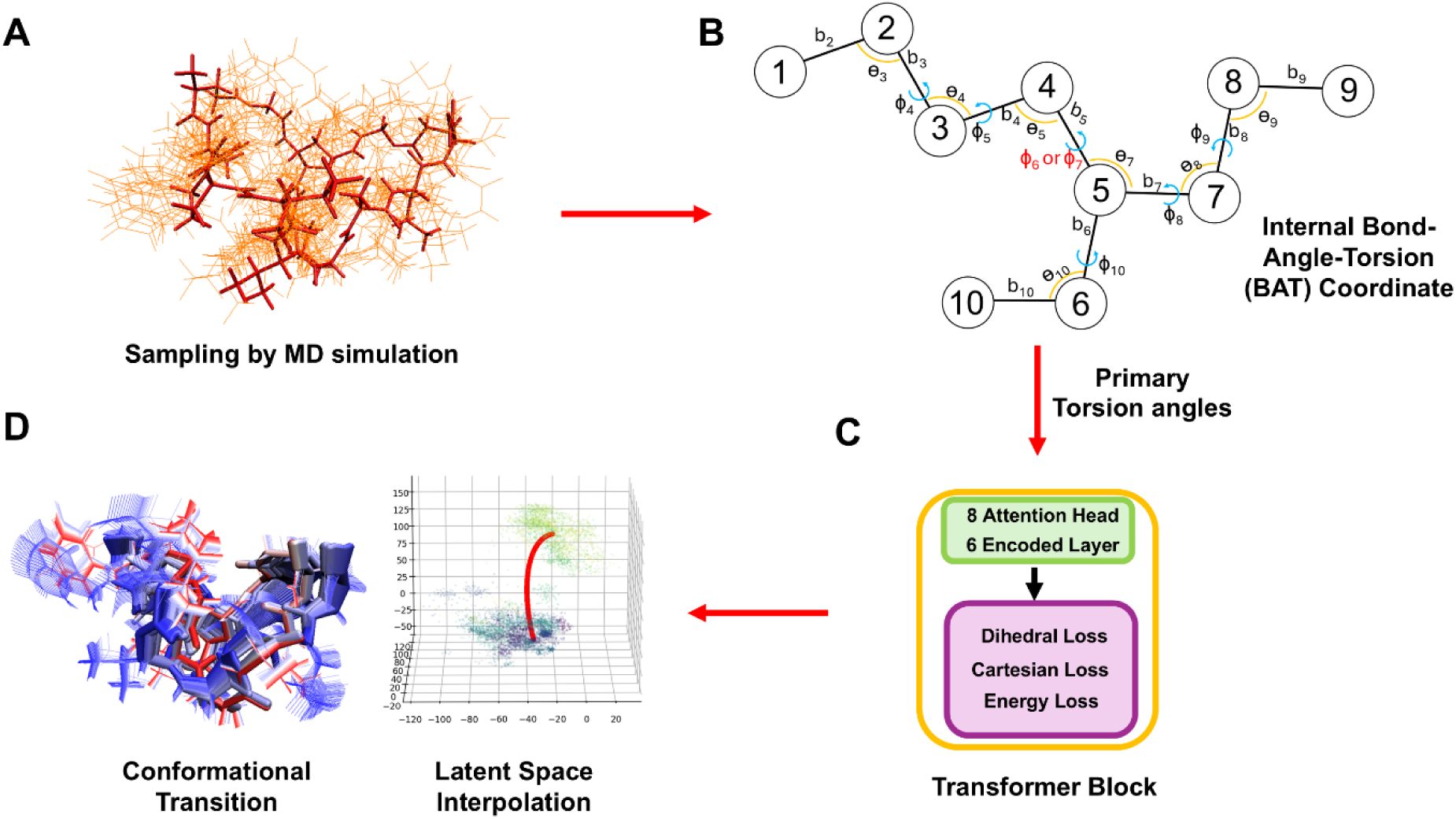
Overview of ICoN-v1. **(A)** The dataset comprises conformational ensembles obtained from molecular dynamics (MD) simulations, with the **(B)** Cartesian coordinates converted to Bond-Angle-Torsion (BAT) coordinates to represent cyclic peptide conformations. The primary torsion angles are extracted as input features for training. The bonds joining atoms 5 with 6 and 5 with 7 can be represented by torsions _6_ and _7_, respectively. _7_ is selected as the primary torsion that leads to conformational changes for atom 7 and _6_ is treated as secondary torsion, which is excluded in the input features. **(C)** Transformer model architecture with 8 attention heads and 6 layers (see details in Methods). **(D)** After training, each cyclic peptide conformation is presented as a single point in latent space. Any point in the latent space can be decoded to an atomistic cyclic peptide conformation, including novel synthetic conformations that are not present in the original MD dataset. Interpolation between different local minima, shown as a red curve in the 3D latent space, reveals conformational transition of the cyclic peptide.

Bonds and angles do not contribute to the change of protein conformation. Torsional rotation dominates the conformational transition, and thus only torsions will be used for training the model. This significantly reduces the number of features from 3N-6 to roughly N-2 while preserving complete structural and dynamic information.

To accurately perform mathematical operation in BAT internal coordinates, we needed to address the discontinuity issue at (±180°) for torsion rotation. For instance, dihedral rotation from 179° to −179° is actually a 2° shift in angular space. However, the basic arithmetic operation will result in a large rotation, 179°-(−179°) = 358° ^24^. To address the discontinuity issue, we introduced angle-aware arithmetic by mapping each torsion angle θ on the unit circle: (cos θ, sin θ).

#### Primary Torsion Angles

As illustrated in **Figure 2B**, one primary torsion is selected for each bond in the molecule whose rotation is the most important one to drive conformational changes. For example, the bonds joining atoms 5 with 6 and 5 with 7 in **Figure 2B** can be represented by torsions □_6_ and □_7_, respectively. Note that □_7_ is selected as primary torsion to that leads conformational changes for atoms 6 and 10, and □_6_ is treated as secondary torsion which is excluded in the input features. Assigning primary torsion allow for a more accurate description of dihedral rotation and also recide the number of features from roughly N-2 to (N-2)/3.

### Model Architecture

Our ICoN-v1 model is built on Transformer architecture ^25^. First, the input embedding layer linearly projects each torsion angle, (*sin(θ), cos(θ)*) pair, from 2 to 512 dimensions (**d_model =512**), creating a rich feature space for subsequent processing. A positional encoding mechanism then enhances these embeddings by adding fixed sine/cosine wave patterns that enable the model to recognize the structural significance of positional differences within the molecular chain. A positional encoding is added to these embeddings using fixed sine/cosine functions, enabling the model to distinguish torsions by their position along the molecular chain and to learn structure-dependent correlations that arise from molecular topology and connectivity. The encoder consists of eight multi-head self-attention blocks (**n_heads= 8**) enabling each torsion angle to be represented as a function of its correlations with all other torsions. Six layers (**n_layers =6**) were used for both the encoder and decoder with **GELU** activation function. The encoder output is projected into a compact three-dimensional latent space (**d_latent = 3**). The decoder section mirrors the encoder structure in reverse that regenerates a 512 dimensional embedding for every torsion angle, followed by a final linear layer that converts these rich embeddings back to predicted (*sin(θ), cos(θ)*) pairs.

#### Learning Rate Scheduler

Learning rate scheduler linearly increases the learning rate during the initial five percent of the training process to prevent training instability, followed by a smooth cosine decay trajectory that gradually reduces the rate towards a user-specified minimum value (**Learning rate = 0.0001 in this study**), optimizing convergence behavior.

#### Loss Function

To improve the accuracy of model prediction and efficiently learn physics, four losses: Torsion, Cartesian, Energy, and Interpolation Energy loss are added in multiple stages of the training progress. A simple Mean Square Error (MSE) loss function is used to assess the accuracy of prediction during the training progress by comparing the model input and output.

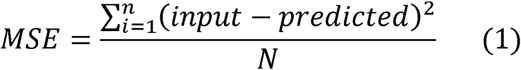

##### Torsion Loss

Torsion loss is defined as the mean squared error (MSE) between the predicted torsion angles and the input torsion angles. During the first training epoch, only the torsion loss is used. Because torsion loss converges quickly, it provides a stable foundation for subsequent training stages.

##### Cartesian Loss

Cartesian loss is defined as the MSE between predicted and the input all-atom Cartesian coordinates. It is used throughout the rest of the training process to ensure the reconstruction accuracy of the model. Computing Cartesian loss is relatively expensive, as full molecular conformations must first be reconstructed from BAT internal coordinates, and this reconstruction is sequential, one atom at a time.

##### Energy Loss

After reconstructing predicted conformations, we compute their potential energy terms, bond, angle, torsion, van der Waals (vdW), and electrostatic (Elec), and the solvation energy term using the Generalized Born (GB) model to avoid unrealistic Elec attraction observed in vacuum. Unlike torsion or Cartesian loss, the energy itself is used as the loss, which explicitly drives the network toward energy minima. This loss penalizes unphysical clashes and stretching, encouraging the model to learn conformations that are physically realistic. Since energy evaluation must be performed for every predicted conformation at each iteration and epoch, this step is computationally expensive. Moreover, because energy values are on a different scale than torsion or Cartesian losses, we apply an adaptive weighting scheme to balance their contributions without bias **(see Adaptive Weight section)**.

Interpolated Energy Loss To further enforce physical realism in latent space, we apply random spherical linear interpolation (SLERP) between latent vectors (**see Random Spherical Linear Interpolation (SLERP) section)**. Along each interpolation path, random points are sampled, decoded into conformations, and their corresponding energy terms (bond, angle, torsion, vdW, Elec, GB) are computed Unlike torsion or Cartesian loss, the interpolated energy itself is used as the loss, which explicitly drives the network toward energy minima. This term ensures that the latent space itself encodes physically meaningful transitions between conformations.

#### Random Spherical Linear Interpolation (SLERP)

The training dataset cannot cover the entire space in the latent space. The empty space does not have the reference conformations to compute the cartesian loss. Our model implements random spherical linear interpolation (SLERP) between points in latent space and randomly selects a point on the spherical path shown in equation 2 and 3. The energy of the interpolated conformations is then computed and included in the loss function. In other words, the network will minimize the interpolated loss/energy during the training process. This process serves as regularization as high-energy regions discovered during interpolation provide learning signals that guide the network to avoid unnatural conformational regions, ultimately leading to a more physically realistic latent space representation.

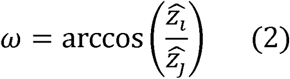

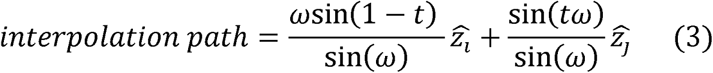

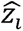 represent the first point and 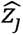 represent the last point in the latent space. *ω* is the angle between 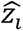 and 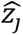. *t* is the parameter to move along the spherical path between 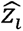 and 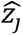 where 0 ≤ *t* ≤ 1.

#### Adaptive Weight

An adaptive weight function to dynamically adjust the contribution of the energy loss terms is included to ensure stable training progress and high geometric accuracy. This function computes the appropriate energy weight by considering the RMSD of the reconstruction and the current training progress. Specifically, the function derives scaling factors for bonded and non-bonded energy terms, ensuring that energy weight is proportionally adjusted based on structural accuracy and if the RMSD of the reconstruction increases, the energy terms are scaled down. This ensures that the conformation ensemble is not overly minimized to similar conformations. Furthermore, the function incorporates a gradual introduction mechanism for energy terms, activating them only after 20% of training has been completed and then linearly increasing their weight until they reach full influence at 70% of the training process. This careful balancing ensures that energy terms do not dominate during early training phases when structural representation is still being established and allows for proper integration of geometric accuracy with energetic constraints as training progresses.

#### Latent Space Sampling through Minimum Energy Pathway (MEP)

After training, each conformation of the MD simulation was encoded as each unique data point in the 3D latent space (**Figure S1A**). Interpolation between two data points in 3D latent space is nontrivial. Because intermediate data points are lacking, there is no constraint to guide a physically meaningful transition, and interpolation through this empty region could yield infinitely many arbitrary trajectories. A realistic conformational transition, however, should follow the MEP, which represents the most favorable route between the two states. To address this limitation, we applied PCA to project the 3D latent representations onto a 2D principal component space (**Figure S1B**). We then computed the probability density distribution of the 2D data and visualized it as a contour map, which revealed local minimum and the corresponding energy gradients (**Figure S1C**). From the 2D probability contour map, we identified two representative conformations, MD1 and MD2, each located at a distinct local minimum of the 2D contour plot. Then, interpolation was subsequently performed along the energy gradient connecting MD1 and MD2 within the 2D probability landscape, providing an approximation of the MEP (**Figure S1D**). Finally, we can map the 2D MEP back to the closest point in 3D latent space and apply polynomial fit applied to create a smooth 3D MEP path (**Figure S1E**). The 3D MEP is reconstructed back to the 3D conformation through decoder. The smooth conformational transition pathway can be obtained and analyzed to understand the mechanism in full atomistic detail (**Figure S1F**).

#### Post Training Analysis

Each interpolated trajectory undergoes a brief energy minimization to remove unfavorable bond stretching and angle distortions which can otherwise dominate the potential energy landscape even under small distortion. We perform 1,000 steps of minimization using the sander module in the AMBER24 suite ^26^, consisting of 800 steps of steepest descent followed by 200 steps of conjugate gradient minimization.

After minimization, the potential energies of the interpolated trajectories are computed using the Molecular Mechanics/Poisson–Boltzmann Surface Area (MM/PBSA) framework in the AMBER package ^27^. Here we only report MM energy terms. A small number of explicit water molecules may reside within cyclic peptides and introduce inaccuracies in PB calculations. We therefore excluded PB from our conformational energy comparisons. Hydrogen bond analysis is performed using VMD ^28^, and torsion angle plots are generated and analyzed using T-analyst ^29^ and in-house scripts.

## Results and Discussion

### Overview of ICoN-v1 network architecture and training

To ensure the high quality of the training dataset, we used trajectories generated by all-atom MD simulations with explicit solvent model to train our ICoN-v1 model. We first converted Cartesians to internal BAT coordinates. This representation removes 6 external rotational and translation degrees of freedoms (DOFs) to yield 3N-6 DOFs (i.e., N-1 bonds, N-2 angles and N-3 torsions), where N is number of atoms. Because rotating a few atoms (i.e., methyl rotation) may be described by a single primary torsion, we used “primary” torsions as the input features for model training (see details in Methods). Instead of using all output frames from MD runs, we performed quick minimizations for all frames and removed the repeating conformation with a 1.5-Å cutoff root mean square deviation (RMSD) for heavy atoms. This allows the model to efficiently learn from the distinct conformations.

ICoN-v1 is built on a Transformer architecture composed of six layers with eight attention heads per layer, enabling the model to efficiently learn correlations among torsional motions. During training, the encoder maps each conformation to a unique point in a three-dimensional (3D) latent space, and the decoder reconstructs the corresponding full conformation from this latent representation. Each molecular conformation was encoded by the neural network into a 3D latent space (**Figure 2**), with each point corresponding to a distinct conformation. Novel synthetic conformations during conformational transitions are obtained by interpolating between two latent points (i.e., two conformations with different ring puckers) and decoding the resulting intermediate latent points back into molecular structures. This procedure yields smooth and physically interpretable transition pathways between selected conformations **(Figure 2)**.

### Exploring transient conformations with latent sampling along minimum energy path

We first investigated cyclo-(<u>T</u>VGGVG) and cyclo-(<u>V</u>VGGVG) (**Figure 1A**) to demonstrate how to use ICoN-v1 to yield smooth conformational transitions and explain how mutating THR1 to VAL1 disrupted the highly packed intramolecular H-bond observed in THR1. The training data were obtained from NMR and MD simulations by the Lin group ^21^. After encoding each conformation into the latent space, the dataset can be grouped by two well-defined clusters for cyclo-(<u>T</u>VGGVG): Cluster 1 (purple circle) represents a state sampled only in MD simulations, and Cluster 2 (green circle) the dominant conformation observed in both MD and NMR (**Figure 3A**). The sparse region between these two clusters contains few or no datapoints, so the intermediates are absent in the reported conformations.

**Figure 3.**
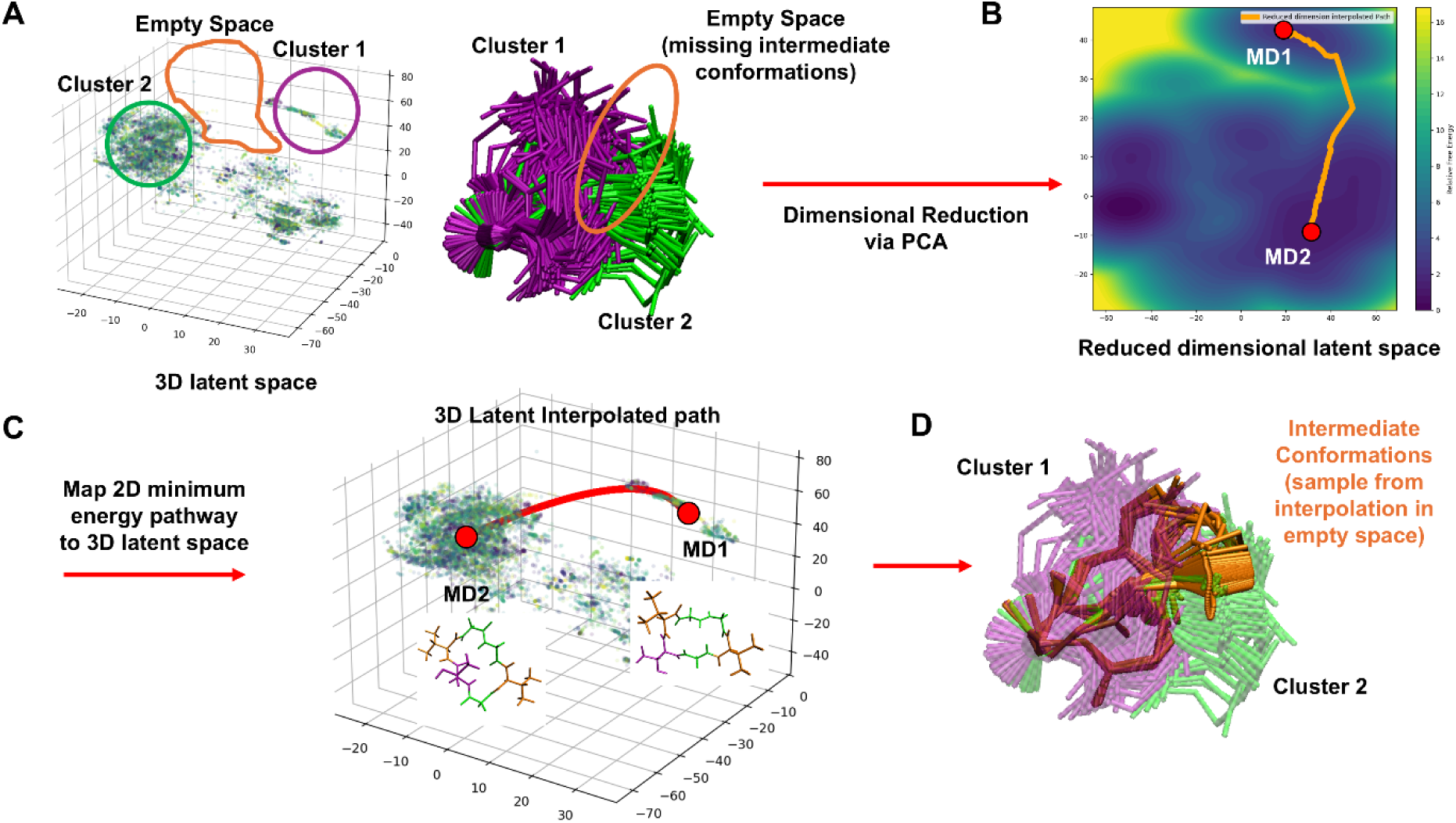
Minimum-energy pathway (MEP)-guided conformational transition of cyclo-(<u>T</u>VGGVG). (**A)** (Left panel) 3D latent space of cyclo-(<u>T</u>VGGVG). Each dot represents different frames of the molecular dynamics (MD) conformation used as training data encoded in the latent space. The green circle denotes Cluster 1, the purple circle denotes Cluster 2, and the orange outline marks the gap where intermediate conformations are missing. (Right panel) Overlaying 100 conformations of Cluster 1 (purple) and Cluster 2 (green). (**B)** A 2D probability contour map generated by projecting latent space datapoints using principal component analysis, with two local minima, MD1 and MD2, selected as interpolation endpoints. The orange path indicates the MEP connecting MD1 and MD2. **(C)** The 3D MEP (orange path in **B**) is mapped back onto the 3D latent space, producing the red curve that guides interpolation. **(D)** Overlaying 100 synthetic conformations (orange) generated along the red curve in **C** reveals the missing intermediate conformations between Cluster 1 (purple) and Cluster 2 (green).

To sample conformational transition, one can interpolate points in the latent space constructed by ICoN-v1, as demonstrated in **Figure 2D**. However, arbitrarily interpolating the two clusters does not guarantee a thermodynamically favorable transition pathway. We instead determined a minimum-energy pathway (MEP) in the latent space that follows the underlying energy gradient and connects two local energy minima through the most probable route. Constructing an MEP directly in the 3D latent space is numerically challenging because of the difficulty of building a well-resolved probability contour map in 3D. We projected the latent space onto two principal components, reducing it to 2 dimensions (2D), which allows for easier construction of probability contour maps (see Methods and **Figure S1** for details). After selecting two local minima, MD1 and MD2, which are also conformations in Cluster 1 and Cluster 2 sampled by MD runs, a 2D MEP path was generated following the energy gradient, which can be easily visualized in the 2D space (**Figure 3B**). The 2D MEP path was then mapped back to the 3D latent space and polynomial fitting was used to reconstruct the 3D MEP for performing a mathematically continuous interpolation. Notably, except for MD1 and MD2, other points in the 3D MEP were not present in the MD simulations (**Figure 3C**). Novel synthetic conformations generated by interpolating points along this 3D MEP path in latent space produced a smooth and continuous conformational transition from MD1 to MD2, as shown by the orange conformations in **Figure 3D**.

Of note, the proposed MEP represents only one possible transition pathway. In principle, cyclic peptides may use substantially different conformational pathways, especially when they rearrange conformations to adapt to distinct environments. For example, during membrane permeation, a cyclic peptide can undergo a transition route that differs substantially from conformational transition pathways observed in an aqueous environment, exhibiting the shift from water to the nonpolar membrane interior. To further explore transitions driven by environmental changes, one could intentionally interpolate energy paths that pass through high-energy regions to probe rare or transient conformational transitions, such as those occurring during membrane permeation. Moreover, thorough conformational sampling may be achieved by systematically, or even exhaustively, exploring transition pathways between datapoints in latent space. Analyzing these paths may uncover distinct sets of concerted torsional rotations and their energy landscapes, which help to identify molecular mechanisms, quantify energy barriers and estimate kinetic properties.

### Revealing concerted torsional rotations during conformational transition of hexacyclic peptides

The MEP-guided interpolation revealed a stepwise conformational transition between two local minima MD1 and MD2, for both hexacyclic peptides cyclo-(<u>T</u>VGGVG) and cyclo-(<u>V</u>VGGVG) (**Figure 4-A1 and -B1**). The potential energies computed along the interpolated curve indicate that after crossing an energy barrier, cyclo-(<u>T</u>VGGVG) transitions from the minimum MD1 to intermediate state T1, then proceeds to another intermediate state T2 before reaching the global minimum MD2 sampled by both NMR and MD (**Figure 4-A2**). Because no original datapoints represent either the T1 or T2 state, we selected frames 1800 and 4000 to represent T1 and T2 conformations, respectively (**Figure 5**). Analyzing their torsional rotations revealed that concerted rotating of GLY3/6 (Phi) and THR1 (Psi) led MD1 to T1, which slightly altered the ring pucker and lost the intramolecular H-bond. Subsequently, GLY6 (Phi) switched again, accompanied by rotation of VAL5 (Psi) to move T1 to T2 to strengthen the polar attraction and for the H-bond to reappear. A new set of concerted torsion rotations, THR1 (Psi), GLY3 (Phi), GLY4 (Phi), VAL5 (Psi), and GLY6 (Phi) shown in **Figure 5-MD2** and detailed in **Figure S2**, changed the ring pucker from T2 to more folded MD2.

**Figure 4.**
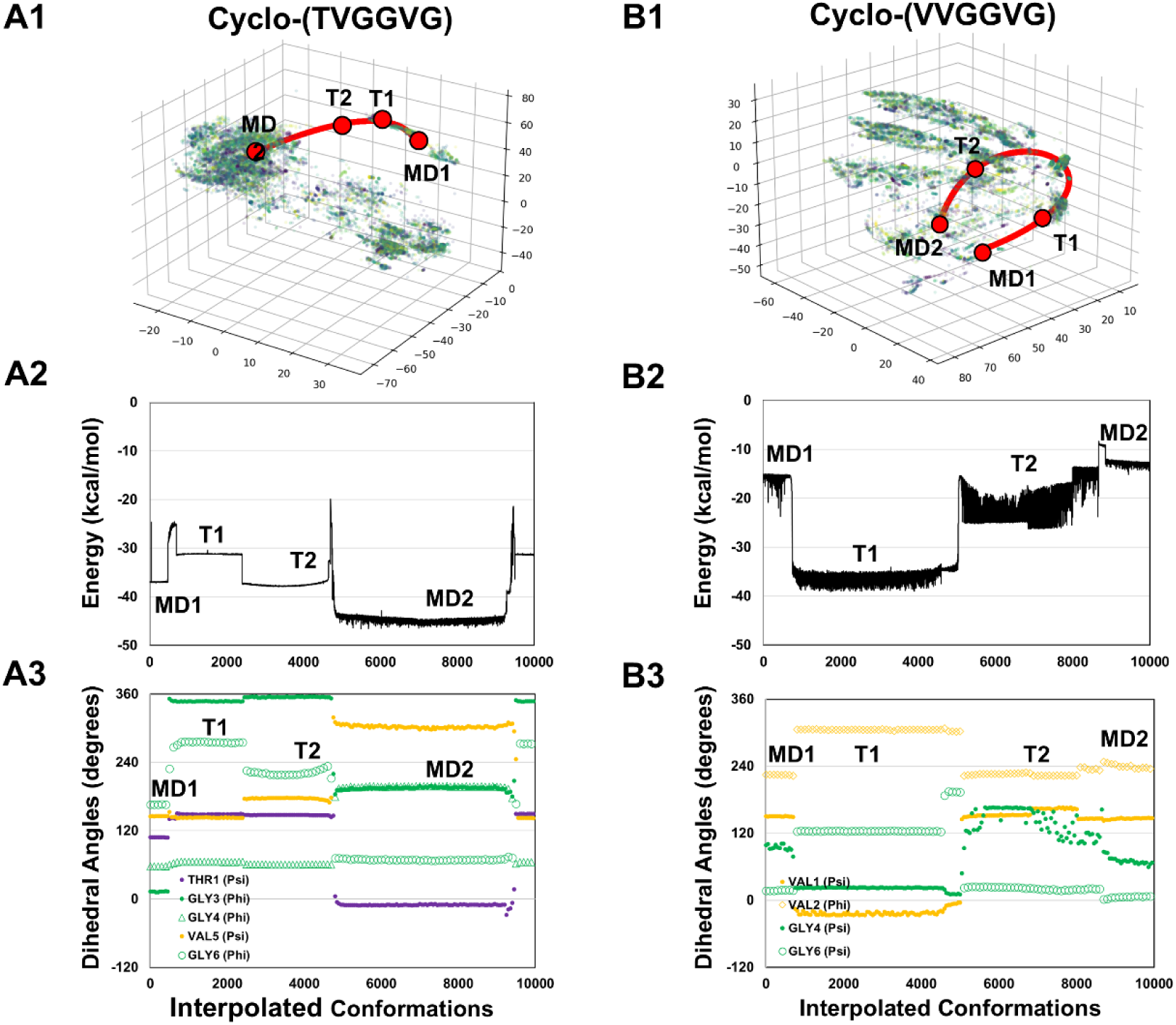
Conformational transition pathway of cyclo-(<u>T</u>VGGVG) and cyclo-(<u>V</u>VGGVG). **(A1 and B1)** Training datapoints in the 3D latent space: different frames of molecular dynamics (MD) frames encoded in latent space are represented in different colors. The minimum-energy pathway (MEP)-guided interpolation curve (red curve), with two end datapoints, MD1 and MD2, and local minima, T1 and T2, are the key transition steps. (**A2 and B2**) Computed potential energy for conformations along the interpolation curve (red line in **A1 and B1**). (**A3 and B3**) Changes in the major torsion rotations of the interpolation curve (red line in **A1 and B1)** contribute to the conformational transition.

**Figure 5.**
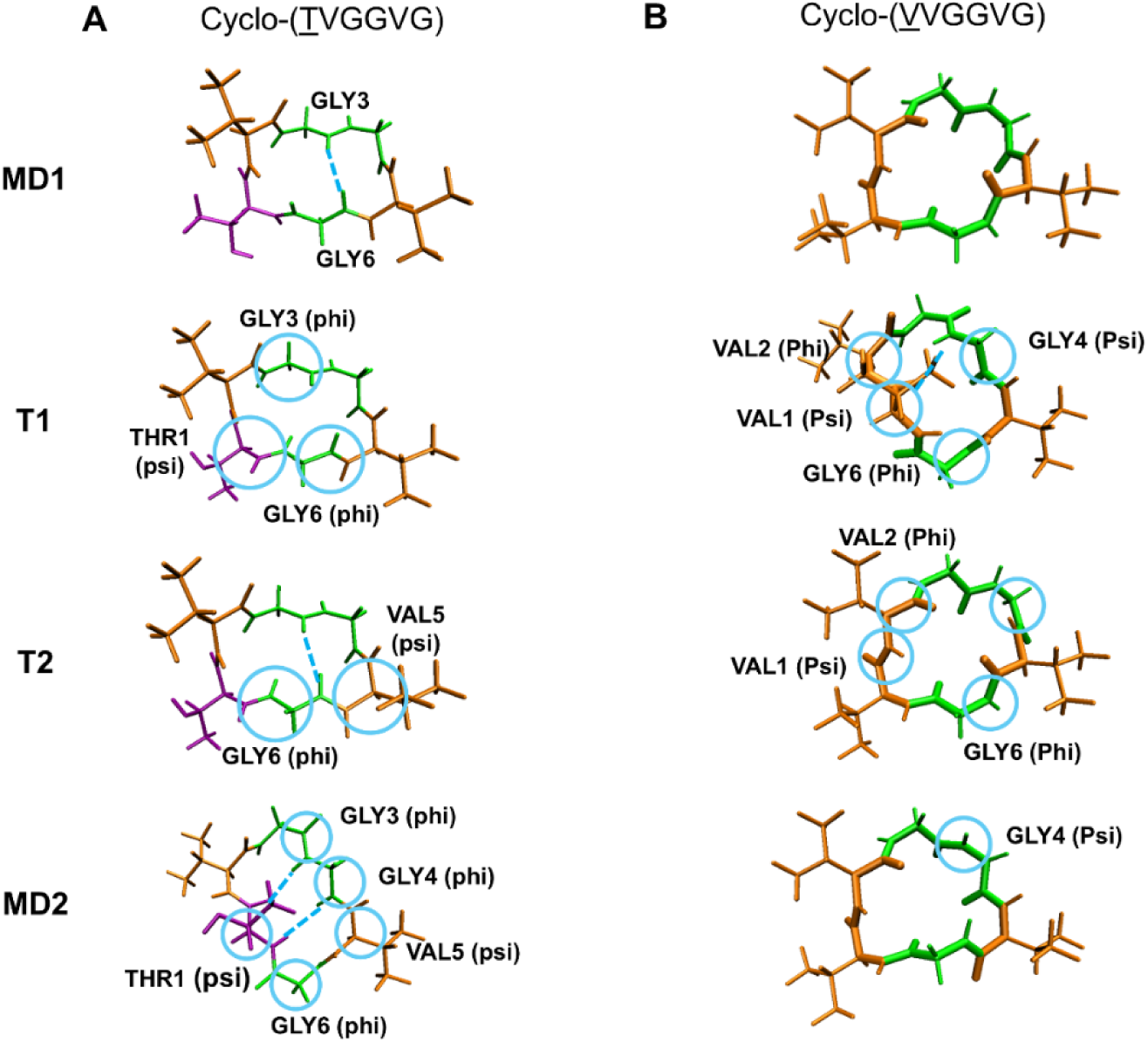
Representative conformations of minimum-energy pathway (MEP)-guided interpolation curve (red line of **Figures 4A1** and **4B1**). Four conformations, MD1, T1, T2, and MD2, are shown in (A) cyclo-(<u>T</u>VGGVG) and (B) cyclo-(<u>V</u>VGGVG). The hydrogen bonds that exist in the conformations of the low-energy minima, MD1 and MD2 of cyclo-(<u>T</u>VGGVG) and T1 of cyclo-(<u>V</u>VGGVG), are labeled in blue dashed lines. The important torsion rotations are circled in blue.

Starting from an arbitrarily selected local minima MD1 shown in **Figure 4-B1**, MEP-guided interpolation led cyclo-(<u>V</u>VGGVG) to the global energy minimum at T1 and then to cross the energy barrier to T2 and finally reach another local minimum at MD2. Of note, the conformation of MD1 and T2 were similar, and our model showed that similar sets of dihedrals are used when transitioning from MD1 to T1 and from T1 to T2. The only difference was that GLY4 (psi) rotated from 100 to 0 degrees from MD1 to T1 (**Figure 4-B2 and 5B**) but from 0 to 140 degrees from T1 to T2 (**Figure S3**). This finding highlights the importance of GLY4 (psi) to pre-organize the conformations to MD2 if all dihedral rotations remain the same, then we can expect that T2 will likely return to MD1. This observation also highlights the ability of our model to capture subtle differences in the latent space.

In comparing the two hexacyclic peptides, cyclo-(<u>T</u>VGGVG) moves its polar sidechain from an outward orientation (T2) to an inward position (MD2), forming intramolecular H-bonds with GLY3 and GLY4 and producing a compact ring conformation by THR1/5 (psi) rotation and GLY3/6 (phi) forming stabilizing H-bonds in the global minimum (**Figure 5A**). In contrast, cyclo-(<u>V</u>VGGVG) reorients the backbone carbonyl to form an H-bond with the GLY4 –NH group, changing the ring from a wide-open MD1 conformation to the more compact global-minimum T1 structure by rotating VAL1/2 (phi) (**Figure 5B-T1**), which is adjacent to the mutation site and serves as a key contributor. Its contribution is despite not directly participating in hydrogen bonding because the steric hindrance from the VAL1 sidechain of cyclo-(<u>V</u>VGGVG) prevents the highly packed H-bond contact observed in cyclo-(<u>T</u>VGGVG) (**Figure 5A-MD2**). Such transition pathways are difficult to resolve directly from conventional MD simulations in which transient conformations are short-lived. In contrast, the latent representation learned by ICoN-v1 captured the concerted torsional rotations in different transient states, which emerged naturally from the learned energy landscape, thus providing mechanistic insight beyond what can be inferred from typical MD analysis alone.

### Conformational transitions of stereo-diversified bicyclic peptides

We investigated five bicyclic peptides with the cyclization step connecting residues GLU1, LYS6 and GLU9 to the central linker. The EFPIVK-linker-ALE sequence covalently connects the residues to the C1, C3 and C4 carbons, whereas the EWPHRK-linker-R<u>L</u>E and EWPHRK-linker-RIE sequences connect them to C1, C2 and C4. As a result, we explicitly denote only the C3 or C2 position (i.e., SS_C3_S or RR_C2_R) when describing their stereo structures. Using the MEP-guided interpolation strategy in the 3D latent space, we investigated how linker chirality reshapes conformational transitions and how one residue LEU/ILE substitution contributes to the conformational transition mechanisms to the bicyclic peptides (**Figure 1B**). Our previous study sampled diverse conformations of stereo-diversified MYC-targeting bicyclic peptides using MD simulations; however, how these bicyclic peptides lead to different conformations was not understood ^22^. All bicyclic peptides studied here have various local minima, including those with open and folded conformations. Not unsurprisingly, folded conformations are more stable in an aqueous environment because they can form more intramolecular contacts. To investigate high- to low-energy conformational transition paths for each system, we constructed MEP (**Figure S4**) and selected two endpoints, MD1 (high-energy state) and MD2 (low-energy state), for sampling their structural transitions using interpolation of MEP-guided curve in the latent space (**Figures 6A**). On the basis of computed potential energy (**Figure 6B**) and corresponding torsional rotations along the interpolation curve (**Figure 6C**), we selected intermediate conformations (i.e., T1-T4) to illustrate the structural rearrangements and the mechanisms driving the transitions.

**Figure 6.**
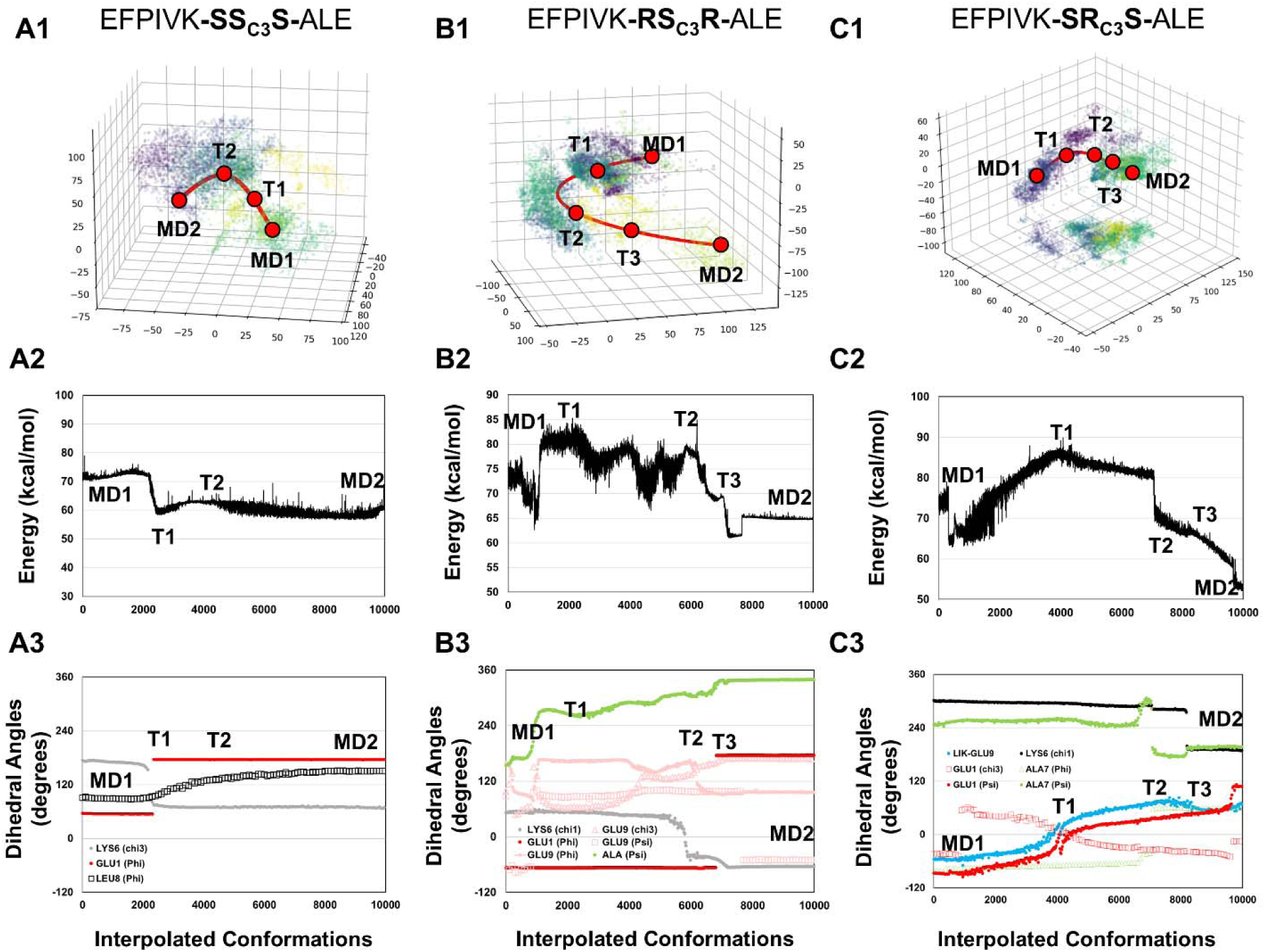
Conformational transition pathways of EFPIVK-SS_C3_S-ALE, EFPIVK-RS_C3_R-ALE, and EFPIVK-SR_C3_S-ALE. **(A1, B1, and C1)** Training datapoints in the 3D latent space; different frames of molecular dynamics (MD) frames encoded in latent space are represented in different colors. The minimum-energy pathway (MEP)-guided interpolation curve (red line), with two end datapoints, MD1 and MD2, and local minima, T1, T2 and T3, are the key transition steps. (**A2, B2 and C2**) Computed potential energy for conformations along the interpolation curve (red line in **A1, B1 and C1**). (**A3, B3, and C3**) Changes in the major torsion rotations of the interpolation curve (red line in **A1, B1, and C1)** contribute to the conformational transition.

#### How do EFPIVK-SS_C3_S-ALE, EFPIVK-RS_C3_R-ALE, and EFPIVK-SR_C3_S-ALE, with the same sequence but different stereo-structures, vary their high-energy (open) to low-energy (folded) transition?

With guidance by the probability contour map (**Figure S4**), we selected an open conformation (high-energy state) for each bicyclic peptide. Not unsurprisingly, folded conformations are more stable (low-energy state) in an aqueous environment because they can bury non-polar sidechains to form more intramolecular contacts while extending polar sidechains to interact with water molecules. To investigate open (high energy) to folded (low energy) conformational transition paths for each system, we constructed an MEP and selected two endpoints, MD1 (open) and MD2 (folded) (**Figure 6-A1 and Figure S4**). Although the side chains of GLU1 and GLU9 were spatially separated for the three peptides with the same sequence, the chirality yielded different open conformations (**Figure 7-MD1**). Different open conformations are anticipated because experiments showed that these bicyclic peptides had various binding affinity to the MYC epitope ^22^. Of note, the S configuration in the chiral center C3, EFPIVK-**SS_C3_S**-ALE and EFPIVK-**RS_C3_R**-ALE, shares highly similar folded conformation to MD2, where PRO3 and ILE4 are in close contact with LYS6 (**Figure S5**). However, their transition pathways were remarkably different. EFPIVK-**SS_C3_S**-ALE follows a straightforward transition pathway, with the potential energy landscape featuring only a single intermediate conformation T1. The transition from MD1 to T1 is initiated by rotations of GLU1 (phi) and LYS6 (chi3) to drive PRO3 and ILE4 toward the linker region and alter the ring pucker, followed by rotation of LEU8 (phi) to pass a small energy barrier (T2) (**Figure 6-A2 and -A3**) to optimize the intramolecular attraction of LEU8 with surrounding residues and reach the global energy minimum, the folded conformation MD2.

**Figure 7.**
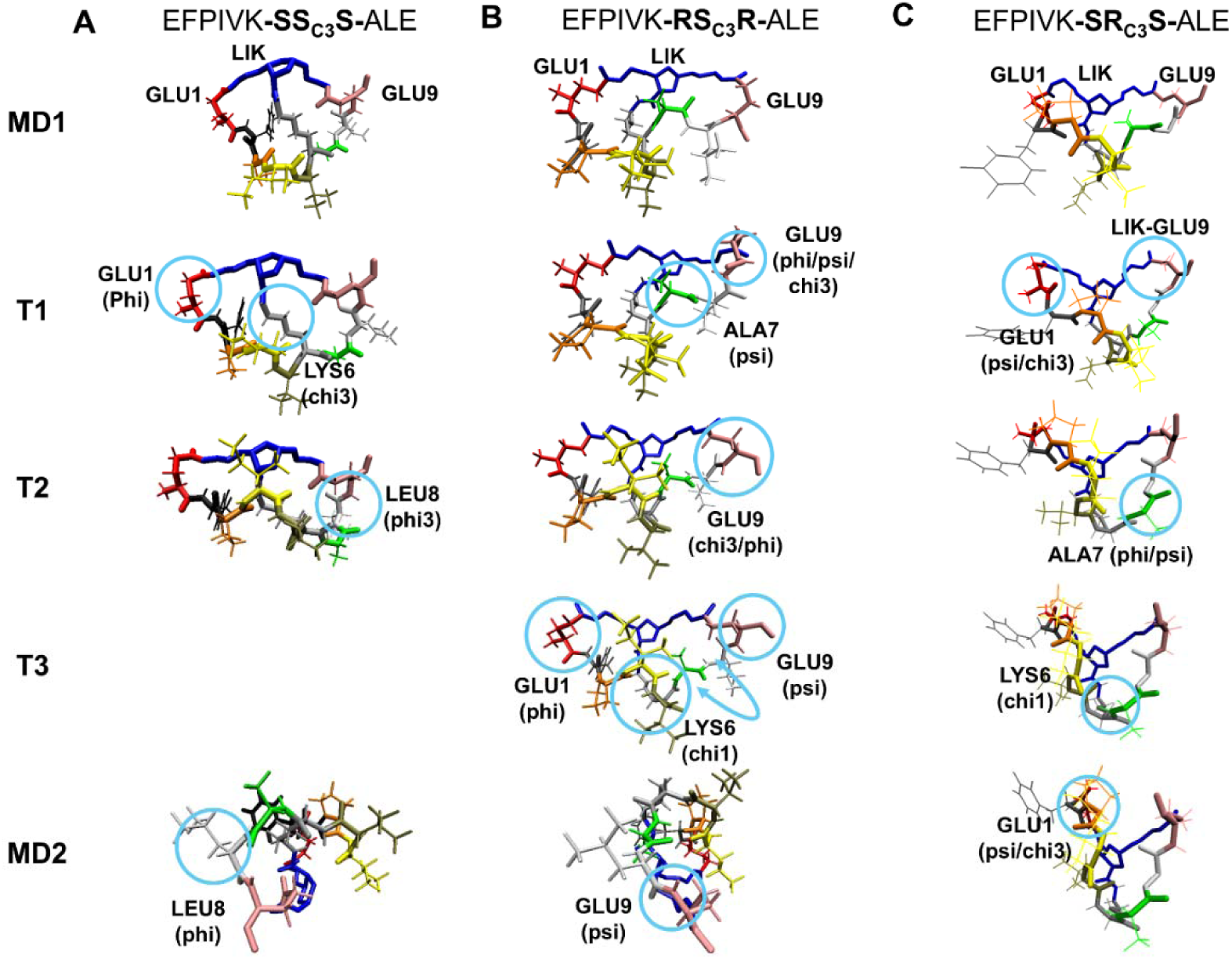
Representative conformations of minimum-energy pathway (MEP)-guided interpolation curve (red lines of **Figure 6-A1, -B1** and -**C1**), MD1, T1, T2, T3, and MD2 of **(A)** EFPIVK-SS_C3_S-ALE, **(B)** EFPIVK-RS_C3_R-ALE, and **(C)** EFPIVK-SR_C3_S-ALE. The important torsion rotations shown in Figure 6**-A3, -B3 and -C3** are circled in blue.

The interpolation curve reveals that EFPIVK-**RS_C3_R**-ALE undergoes an open-to-folded transition via a series of concerted torsional rotations that modulate the ring pucker (**Figure 6-B1**). To characterize this process, we selected three representative intermediate conformations (T1–T3) that capture the key switching events in these concerted torsion rotations. **Figure 6-B2 and -B3** demonstrate that EFPIVK-**RS_C3_R**-ALE follows a distinct set of torsional pathways and traverses a more rugged energy landscape yet ultimately converges to the same folded minimum MD2 conformation observed for EFPIVK-**SS_C3_S**-ALE. ALA7 (Psi) and GLU9 (Phi/Psi/chi3) initiate the conformational transition to overcome ∼20 kcal/mol energy change and bring EFPIVK-**RS_C3_R**-ALE to a local energy barrier (T1); the two torsions continue rotating non-linearly to guide the molecule visiting a few local minima before reaching another energy barrier (T2). To open space for PRO3/ILE4/VAL5 to move inward and interact with the linker, GLU1 (phi), LYS6 (chi3) and GLU9 (phi) on opposite sides of the ring rotate coordinately to reach the local minimum T3, followed by adjusting GLU9 (Psi) to complete the transition to MD2 (**Figure 7B**).

With an R configuration in the chiral center C3, EFPIVK-**SR_C3_S**-ALE has a distinct folding state of MD2, with GLU1 and GLU9 forming close contacts (**Figure 7C-MD2**). As for EFPIVK-**SS_C3_S**-ALE, GLU1 initiates the conformational transition. However, in **Figure 6-C2 and -C3**, EFPIVK-**SR_C3_S**-ALE uses a different set of torsions, GLU1 (psi/chi3) and linker-GLU9, to overcome ∼20 kcal/mol energy and reach a local barrier T1. The additional rotation of ALA7 (Phi/Psi) then reorganizes the backbone to form T2, thus enabling the subsequent rotation of LYS6 (chi1) for the T2 to T3 transition. A final adjustment of GLU1 (psi and chi3) completes the folding process and stabilizes GLU1–GLU9 contacts (MD2) (**Figure 7C**).

Residues GLU1, LYS6, and GLU9 directly connect to the linker region, so rotating backbone phi or psi angles of these residues contributes substantially to changing the ring conformations. However, different stereo-structures use distinct pathways, each identified by specific combinations of torsions in these three residues, coupled with coordinated rotations of torsions in ALA7 or LEU8 to lead to different intermediate conformations and finally achieve an open to folded conformational transition. Although ALA7 does not directly attach to the linker, its backbone rotation can notably fluctuate the ring pucker and largely change the energy by ∼20 kcal/mol (**Figure 6-B2** and **-C2**). Of note, torsional rotations in PRO3 and ILE4 were not observed for all three bicyclic peptides with the EFPIVK-linker-ALE sequence. Instead, motions from nearby residues resulted in strengthening interactions between PRO3/ILE4 with other residues (i.e., PRO3-LEU8 or ILE4-linker shown in **Figure S6**). These interactions lowered the potential energy by ∼10 kcal/mol, which suggests their mechanistic importance in stabilizing the conformations and driving the system toward the folded global minimum. Mutating PRO3 or ILE4 to a bulkier residue such as TYR or TRP introduced substantial steric hindrance, which could disrupt the formation of folded conformations and shift the energy landscape toward more opened conformations. Depending on the application, the open conformations may improve binding to their protein target but potentially impair the membrane permeability of the bicyclic peptide.

#### How does a single point mutation, from EWPHRK-RR_C2_R-RLE to EWPHRK-RR_C2_R-RIE, alter the conformational transitions from high to low energy states?

Unlike proteins, in which the highly similar residues LEU and ILE usually yield nearly identical structures, mutating LEU to ILE in EWPHRK-RR_C2_R-R<u>L/I</u>E can completely reshape the conformational ensemble. In contrast to the EFPIVK-linker-ALE bicyclic peptides, which exhibit diverse potential energy landscapes during transitions from high to low energy states, the EWPHRK-RR_C2_R-R<u>L/I</u>E bicyclic peptides follow a smoother downhill energy landscape with no major energy barriers (**Figure 8-A2** and **-B2 and SI Video**). Moreover, both bicyclic peptides share the same high-energy folded conformation (MD1) and a similar open intermediate (T2). In MD1, GLU1 and GLU9 are far apart and sidechains of ARG5 and ARG7 rotate inward to interact with TRP2 and HIS4 to form cation-π interactions to stabilize the folded conformation. However, positioning both positively charged ARG5 and ARG7 in close proximity introduces unfavorable electrostatic repulsion, thus resulting in a high-energy folded conformation. As a result, a natural first step in moving the energy landscape downhill is to relieve the ARG5-ARG7 repulsion by changing the ring pucker. Of note, bicyclic peptides with LEU8 or ILE8 substitutions use different strategies to alter the ring, which gradually leads to two distinct low-energy folded conformations (MD2).

**Figure 8.**
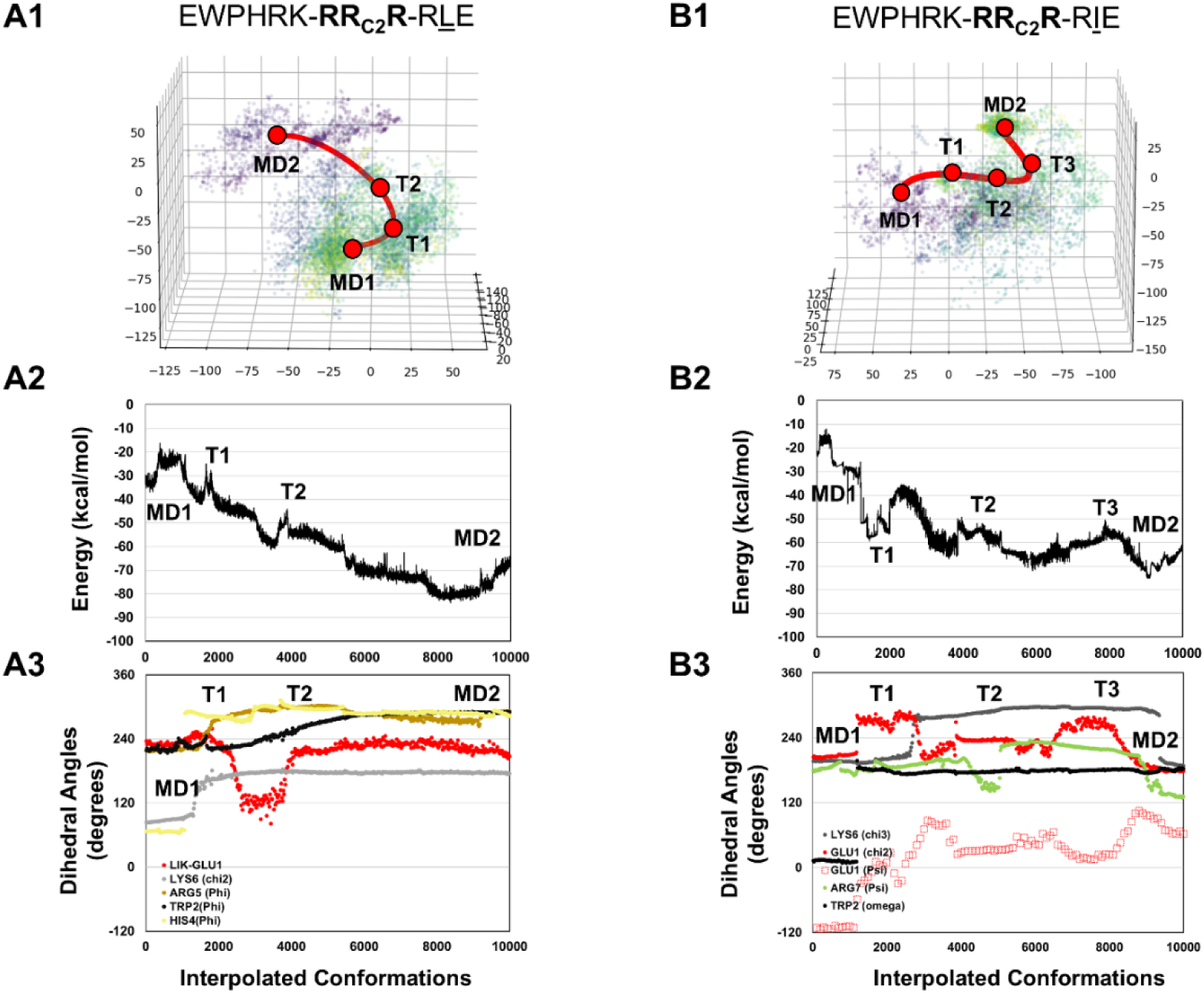
Conformational transition pathways of EWPHRK-RR_C2_R-R<u>L</u>E and EWPHRK-RR_C2_R-RIE. **(A1 and B1)** Training datapoints in the 3D latent space, with different frames of MD frames encoded in latent space represented in different colors. The minimum-energy pathway (MEP)-guided interpolation curve (red line), with two end datapoints, MD1 and MD2, and local minimum, T1, T2 and T3, which highlight the key transition steps. (**A2 and B2**) Computed potential energy for conformations along the interpolation curve (red line in **A1 and B1**). (**A3 and B3**) Changes in the major torsion rotations of the interpolation cure (red line in **A1 and B1)** contribute to the conformational transition.

Beginning from the folded MD1 conformation, outward rotations of TRP2 (phi), HIS4 (phi), and ARG5 (phi) in EWPHRK-RR_C2_R-R<u>L</u>E (**Figure 8-A3**) drive the system into the T1 intermediate state (**Figure 9A-T1**). The concerted rotations then shift torsions HIS4 (phi) and LYS6 (chi2) to guide ARG5 and ARG7 to move outward to produce an open conformation T2. Continued non-linear rotations of TRP2 (phi), ARG5 (phi) and GLU1-linker torsions further refold the ring to another low-energy folded conformation (MD2) (**Figure 9A-MD2**). In EWPHRK-RR_C2_R-RIE, the peptide uniquely rotates the TRP2 omega bond and GLU1 (psi/chi2) to disrupt the cation-π interactions (**Figure 8-B3**) between TRP2/HIS4 and ARG5/ARG7 appearing in MD1, thereby slightly altering the ring pucker (T1) for subsequent rotation of ARG7 to open the ring (T2). The transition from T2 to T3 involves rotations in GLU1 (psi/chi2), with an open conformation, followed by varying degrees of coordinated LYS6 (chi3) and GLU1 (psi and chi2) and ARG7 (psi) rotations to finally form contact between PRO3-GLU9 and HIS4-ILE8, thereby leading to another final folded, low-energy conformation (MD2) (**Figure 9B**).

**Figure 9.**
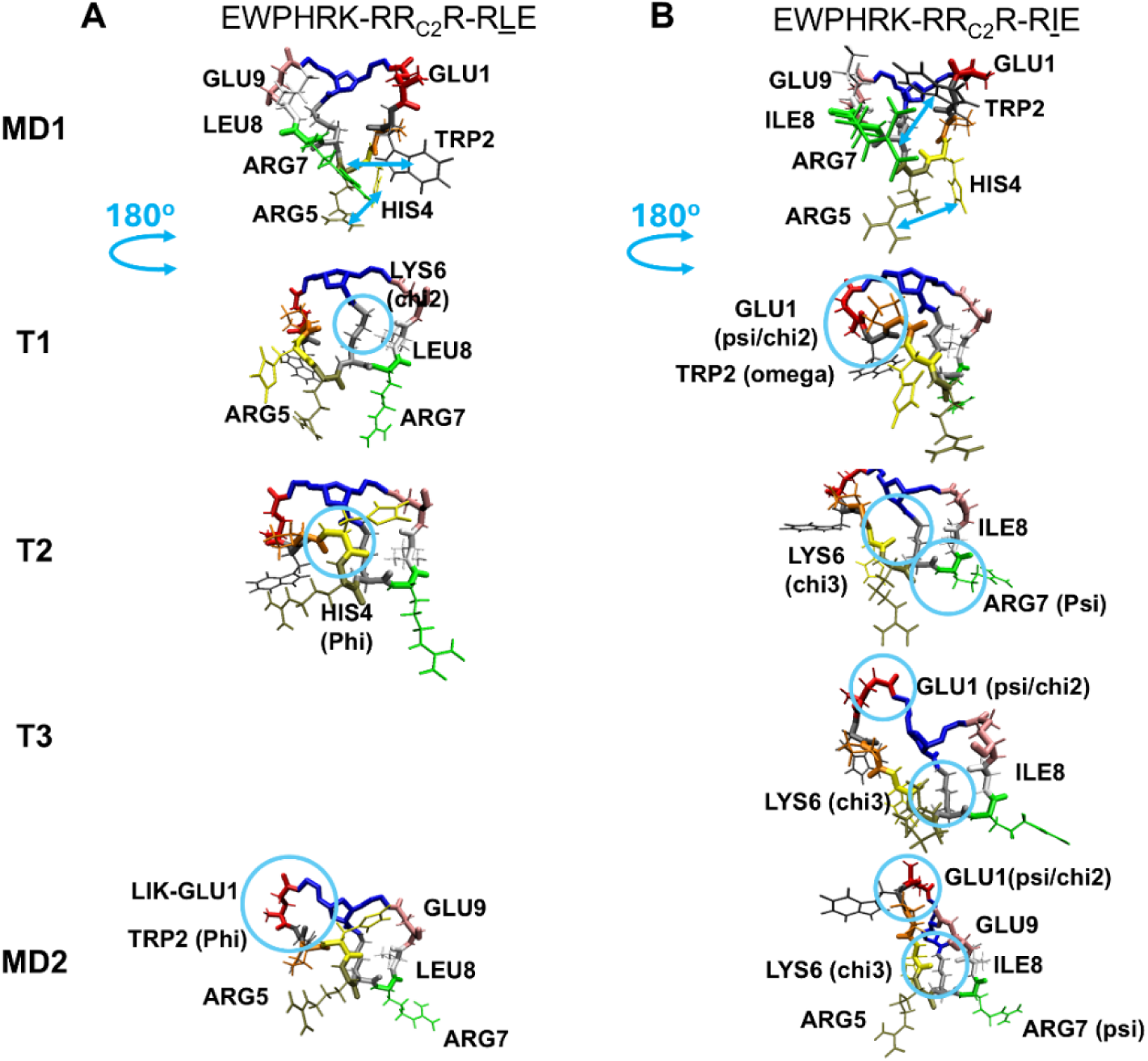
Representative conformations of minimum-energy pathway (MEP)-guided interpolation curve (red lines of **Figure 8-A1** and -**B1**), MD1, T1, T2, T3, and MD2 of **(A)** EWPHRK-RR_C2_R-R<u>L</u>E and **(B)** EWPHRK-RR_C2_R-RIE. The conformations are rotated 180 degrees (shown in blue arrows) from MD1 to T1 for easier visualization of important torsion rotation. The important torsion rotations shown in **Figure 8-A3** and -**B3** are circled in blue.

Investigating the transition pathways for EWPHRK-RR_C2_R-R<u>L/I</u>E explains how LEU8 or ILE8 substitution affects conformational ensembles. Even small sequence changes in bicyclic peptides are well known to produce often unpredictable and large structural variations ^22^. These systems exhibit highly concerted torsional rotations to lead to different conformations. Even when transitioning from one folded conformation to another, a bicyclic peptide may transiently open the ring and then re-fold again into different folded conformations. As demonstrated in EWPHRK-RR_C2_R-R<u>L/I</u>E, even when our latent space interpolation begins from the same high-energy folded MD1 conformation, the two molecules use a completely different set of torsion rotations (i.e., TRP2(phi)/HIS4(phi)/ARG7(phi) in LEU8 substitution and TRP2(omega)/GLU1(psi/chi2) in ILE8 substitution) to disrupt interactions between ARG5/ARG7 and TRP2/HIS4 and change the ring structure. Of note, EWPHRK-RR_C2_R-RIE uses the peptide bond of TRP, TRP2 (omega), to change the molecular conformation. In structural proteins, cis-trans isomerization of an omega bond typically involves a 15∼20 kcal/mol free energy barrier ^30^ and thus often requires a cis-trans isomerase to lower the free energy barrier for the transition. However, in cyclic peptides, the conformational constraints imposed by cyclization lower the energetic barrier for cis-trans omega rotations, thus enabling access to a broader conformational ensemble than typical proteins. This enhanced flexibility allows cyclic peptides to adopt conformations capable of engaging transients or “undruggable” binding sites on target proteins, which highlights a key advantage of cyclic scaffolds in molecular recognition and design.

After the bicyclic ring opens to reach a similar intermediate T2, the bicyclic peptides with LEU8 or ILE8 again use distinct sets of torsional rotations to guide the bicyclic peptides into different low-energy folded conformations, each corresponding to its own MD2 endpoint. Moving LYS6(chi3) refolds EWPHRK-RR_C2_R-RIE (**Figure 8-B3**), where the linear sidechain of ILE8 can be more flexible to adjust its conformation, thus allowing the molecule to adopt a new low-energy folded conformation. However, the branched sidechain of LEU8 is bulkier spatially to prevent rotations of LYS6 from rotating further for conformational transition. After initial changes of LYS6, torsions of LYS6 remain the same through the afterward transitions (**Figure 8-A3**). Instead, although far from LEU8, torsional rotation of TRP2 (phi) plays a pivotal role in bringing HIS4 to contact with the linker, refold the cyclic peptide, and bring the molecule to a low-energy folded form. By examining their transition pathways, we propose that TRP2 functions as a leading residue that differentiates the conformational transition routes.

By illustrating the diverse conformational transitions governed by varying concerted torsional rotations along the pathways, we gain deeper insight into how bicyclic peptides adapt to different environments. Importantly, such an investigation is only possible with a high-quality DL model that learns the underlying physics and captures these dynamically switching sets of torsional rotations. Methods based on linear combinations of torsions using fixed rotational magnitude are inherently unable to present these highly complex and cooperated motions shown here. The folded conformations of bicyclic peptides can diverge enough to have different peptide–water interactions, which may influence membrane permeability. With EWPHRK-RR_C2_R-RIE used as an example, the close packing of ARG5/ARG7 and TRP2/HIS4 in MD1 buries their polar function groups (**Figure S7-MD1**), thus resulting in a peptide–water electrostatic attraction of - 490.4 kcal/mol. This situation is less favorable than in MD2, where both ARG5/ARG7 extend outward to interact with water (**Figure S7-MD2**), thereby increasing the solvent–solute electrostatic attraction to −555.8 kcal/mol. Nevertheless, MD1 with less favorable interactions with polar environments may become the dominant conformation in nonpolar environments, such as within a lipid membrane or in nonpolar solvent. Here, we demonstrate the chameleonic behavior ^31^ of these bicyclic peptides by studying the conformational transition pathways and the concerted torsion rotation.

## Conclusions

This study uses a DL model, ICoN-v1, to reveal the underlying physics governing conformational transitions in bicyclic peptides. By using primary torsion angles to represent atomic motions and incorporating an energy-guided loss function during training, the model learns physically meaningful conformational landscapes and constructs a 3D latent space that encodes molecular conformational information. The latent space can be viewed as a synthetic configuration space that captures the conformational free-energy profile. Interpolating points between local energy minima enables the discovery of conformational transition paths. Analyses of MEP-guided interpolation further offer mechanistic insights into these transitions.

We show how a single mutation with the same residue property substitution LEU8 to ILE8 can introduce distinct transient states and energy barriers via different TRP2 backbone rotations. Our analysis further reveals that chirality alone does not determine the balance between open and folded conformations. Distal residues, such as ALA7 and LEU8, can contribute up to ∼20 kcal/mol to the potential energy landscape, whereas PRO3 and ILE4 stabilize folded conformations. With hexacyclic peptides, we also demonstrate how a THR-to-VAL mutation disrupts the H-bond network and reshapes the free-energy profile. Our model goes beyond end-state comparisons by reconstructing complete transition pathways and exposing intermediate states, which are unreported in existing MD trajectories yet critically govern the conformational ensemble sampled by MD runs. These findings provide mechanistic insights into cyclic peptide conformational landscapes and identify key residues and dihedral angles that control conformational ensembles and energy barriers. Such information can guide rational cyclic peptide design for various applications.

Because detailed, system-specific understanding is essential for molecular design, ICoN-v1 is not intended to construct a universal model, and a system-dependent 3D latent space should be constructed for each molecular system. However, the training dataset is not limited to MD trajectories and can be trained on conformational ensembles generated by any sampling method. However, like many DL approaches, the model can suffer from domain limitations when generating synthetic conformations far from the training distribution. Incorporating physics-based noise, such as Langevin Dynamic style perturbations, may help expand the accessible conformational space and improve the model’s ability to explore previously unseen regions of the energy landscape.

## Data and Software Availability

The code, pre-trained ICoN-v1 model parameters, and data sets are available on https://github.com/chang-group

## Acknowledgements.

We thank Dr. Min Xue for valuable discussion regarding the bicyclic peptide sequences and their associated experimental results. and. We thank Dr. Yu-Shan Lin for providing both MD and NMR conformations of hexa-cyclic peptides. This study was supported by the US National Institutes of Health (R01GM-109045 to C. C.) and the US National Science Foundation (MCB-2437134 to C.C.).

## Notes

### Competing Interest Statement

The authors have declared no competing interest.

### Summary of Updates

We updated the Title and abstract. We rewrite the introduction to make it more concise and clear. We rewrite part of the method to make it concise.

