## Supporting Information for "Mechanistic Dissection of Conformational Transition of Bicyclic Peptide via Molecular Modeling and Deep Learning"

**MD Simulations and Preparation for Training and Validation Datasets**

MD trajectories of the hexapeptides, cyclo-(TVGGVG) and cyclo-(VVGGVG), are obtain from Lin group ^1^. MD trajectories of all bicyclic peptide are obtained from our previous work ^2^. ff14SB force field was used on protein and gaff2 force field was used on linker ^3,4^. The linker charges were computed using AM1-BCC ^5^. All systems were then solvated with TIP3P water box with the extension of 15Å from the solute edge ^6^. The production runs are performed with 2-fs time steps. The MD trajectories were collected at 1-ps interval which made up 1,000,000 frames for each cyclic peptide.

To construct a clean and robust dataset for ICoN-v1 model training, we processed the data to eliminate noisy sidechain rotations and preserve distinct conformations. First, to mitigate uninformative thermal fluctuation, we performed a quick energy minimization on the entire MD trajectories, consisting of 800 steps of steepest descent followed by 200 steps of conjugate gradient. These steps stabilize freely rotating methyl groups which can otherwise introduce bias; because our model is trained solely on primary dihedral angles, it does not inherently distinguish between uninformative methyl rotations and functionally relevant backbone torsions. Second, to preserve only distinct conformations, we calculated the pairwise RMSD for all structures in each trajectory. Using a heavy-atom RMSD cutoff of 1.5 Å, we filtered out repeating conformations. This ensures the model learns from a diverse set of structures rather than being biased toward a single dominant conformation. Each system contains a different number of redundant conformations; therefore, after removing these redundancies, the number of conformations in each training set varies **(Table S1)**.

After preparing a clean dataset, we select four consecutive data points for the training set followed by one data point for the validation set and repeat this pattern until the end of the refined trajectories. Therefore, 80% of the refined trajectories are used as training and 20% are used as validation **(Table S1)**.

**ICoN-v1 Model Validation by Evaluating the Accuracy of Reconstructed Conformations**

To evaluate reconstruction accuracy, conformations from both the training and validation sets were passed through the pre-trained ICoN-v1 model. The reconstructed BAT coordinates were converted back to atomistic Cartesian coordinates, and RMSDs of backbone and all heavy atoms were computed between the original and reconstructed structures. As shown in **Table S1**, the hexapeptide exhibits ~1 Å RMSD for both backbone and all heavy atoms in both the training and validation sets. The bicyclic peptide shows ~1.5 Å backbone RMSD and ~2.0 Å RMSD for all heavy atoms. Our result shows accurate reconstruction in atomistic details including side chain rotation and the robustness of our model in learning diverse conformations. Notably, incorporation of energy loss informs the model to minimize the energy during the training processes. As a result, the model tends to generate locally minimized sidechain arrangements, which can slightly increase the RMSD as compared with a pair of original and reconstructed conformations. Although training solely on coordinate differences can yield smaller reconstruction RMSDs, the underlying physics may not be learned accurately, especially in regions of latent space without datapoints where the model lacks references for computing their Cartesian coordinates loss (data not shown). In addition, the similarity between training and validation RMSDs indicates that the model does not exhibit overfitting. Importantly, the validation set is excluded from training. However, ICoN‑v1 provides validation‑set evaluation results during training, such as original–reconstructed conformation comparisons, which help users tune hyperparameters more efficiently. Our work highlights the importance of incorporating energy terms during training because it provides guidance for any datapoint within the latent space and ensures that the model learns physics accurately.

**Table S1.** Number of molecular dynamics (MD) conformations used in the ICoN-v1 model for model training and validation, and the reconstruction root mean square deviation (RMSD) for backbone and heavy atoms.

| Peptide | Number of raw MD conformations | Number of confs used to build the DL model. 80% for Training and 20% for Validation. | Reconstruction RMSD (Å) training (backbone/all heavy atoms) | Reconstruction RMSD (Å) validation (backbone/all heavy atoms) |
| --- | --- | --- | --- | --- |
| cyclo-(**T**VGGVG) | 100,000 | 17072 | 0.91/1.18 | 0.91/1.18 |
| cyclo-(**V**VGGVG) | 100,000 | 16431 | 0.95/1.16 | 0.95/1.16 |
| EFPIVK-**SS_C3_S**-ALE | 100,000 | 6031 | 1.51/1.83 | 1.51/1.83 |
| EFPIVK-**SR_C3_S**-ALE | 100,000 | 11544 | 1.44/1.72 | 1.43/1.71 |
| EFPIVK-**RS_C3_R**-ALE | 100,000 | 9076 | 1.53/1.81 | 1.53/1.81 |
| EWPHRK-RR_C2_R-R**L**E | 100,000 | 6448 | 1.45/2.16 | 1.46/2.19 |
| EWPHRK-RR_C2_R-R**I**E | 100,000 | 5452 | 1.59/2.29 | 1.59/2.31 |

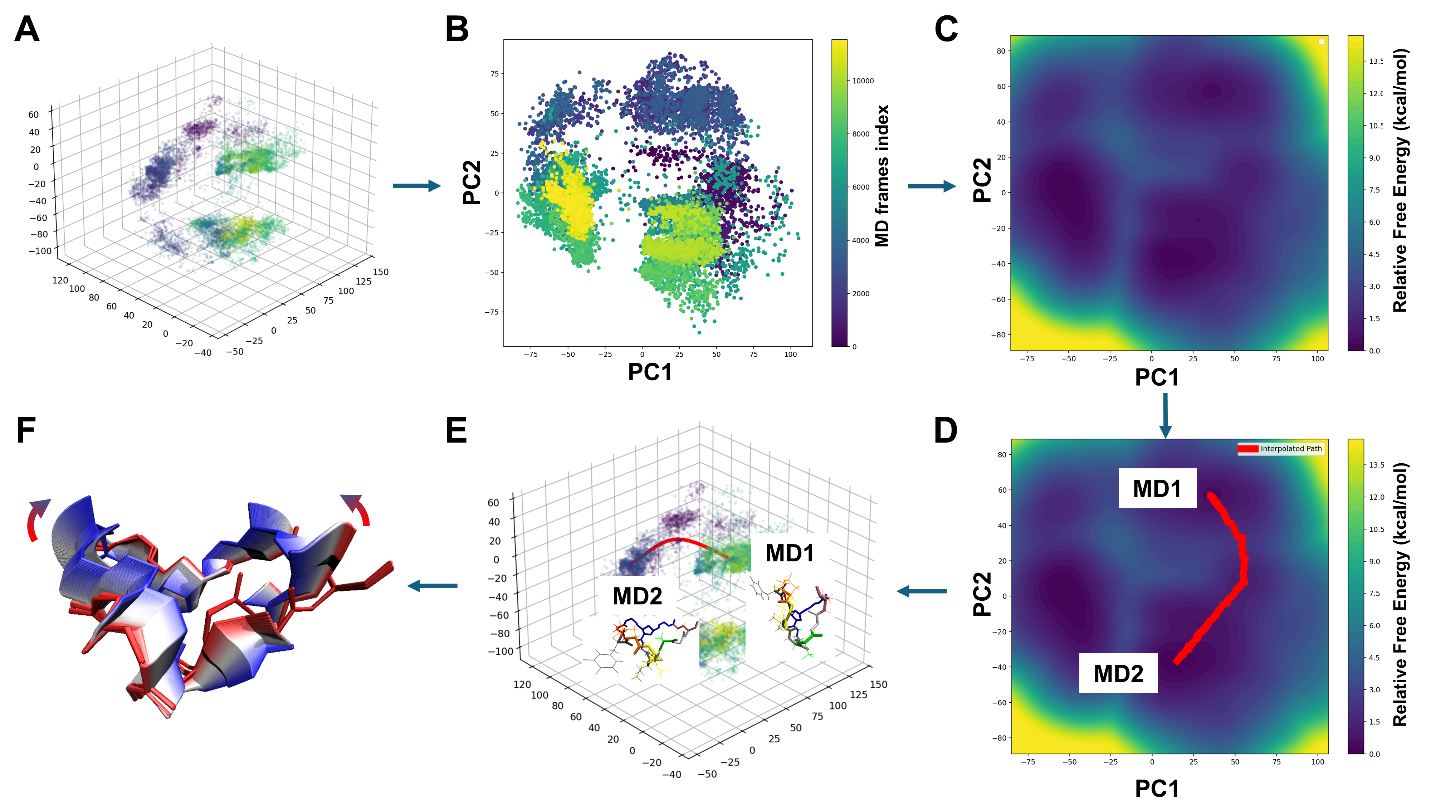

**Figure S1.** Flow chart of interpolation in latent space. **(A)** 3D latent space of ICoN model. Color **(B)** Reduce dimensionality of latent space using PCA. **(C)** 2D probability distribution contour map of PCA **(C)** Local minimums, MD1 and MD2, of the 2D probability distribution contour map are selected as the initial and final conformations of conformational transition pathway. The Minimum Energy Path (MEP) is obtained by following the energy gradient from MD1 to MD2. **(E)** The MEP in 2D space are mapped back to 3D latent space and the polynomial fit is performed to ensure smooth and continuous pathway **(F)** The MEP of the latent space are decoded to cyclic peptide conformations. Overlaying 100 conformations of cyclic peptides of the MEP.

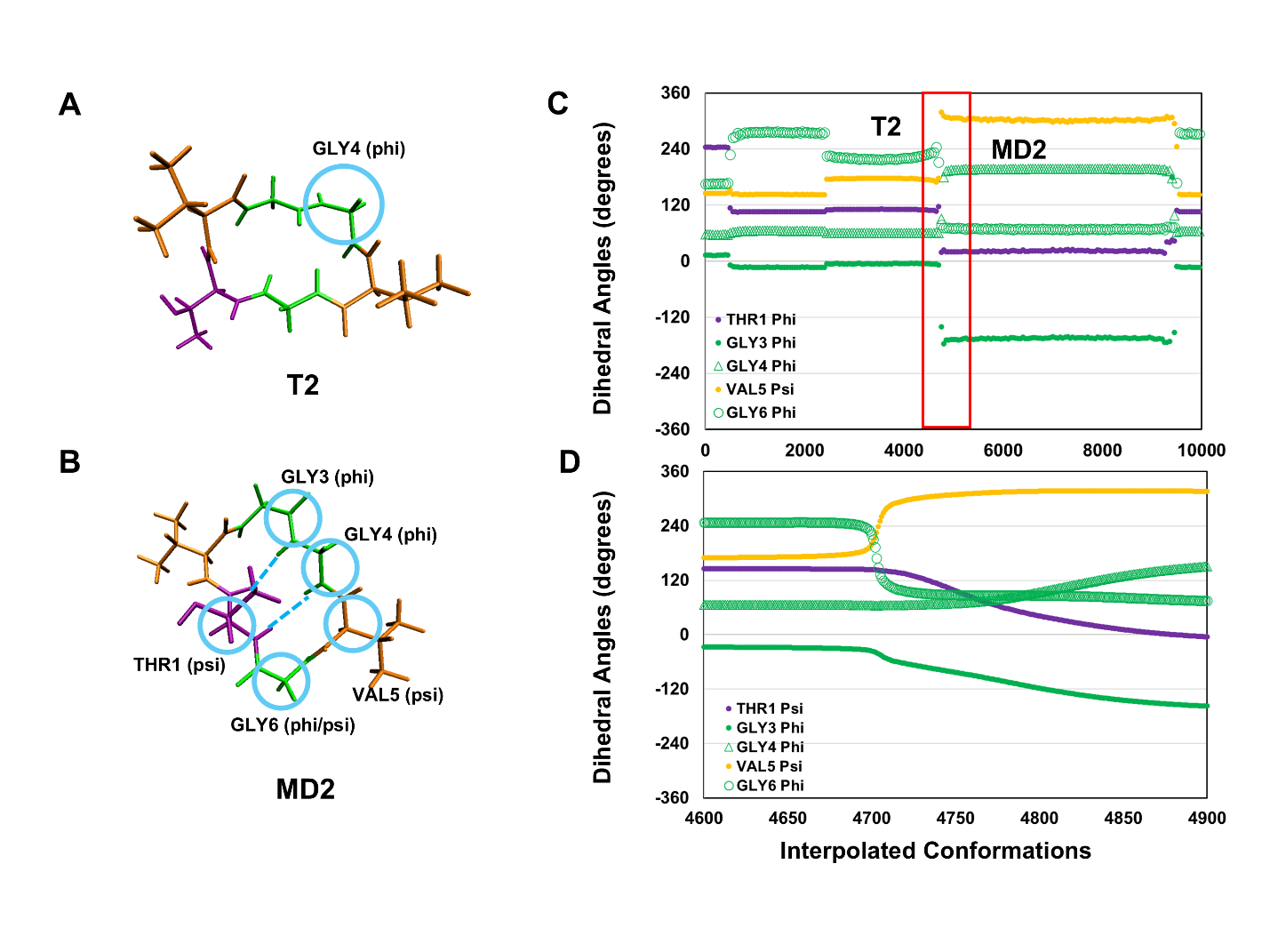

**Figure S2.** Closer observation of cylco-(TVGGVG) conformational transition from T2 to MD2. **(A)** T2 of cyclo-(TVGGVG) from **Figure 4** **(B)** MD2 of cyclo-(TVGGVG) from **Figure 4** **(C)** Change in torsion angle over the interpolation pathway **(D)** Zoom in on the torsion angles during the transition between high energy state (T2) to lowest energy state (MD2).

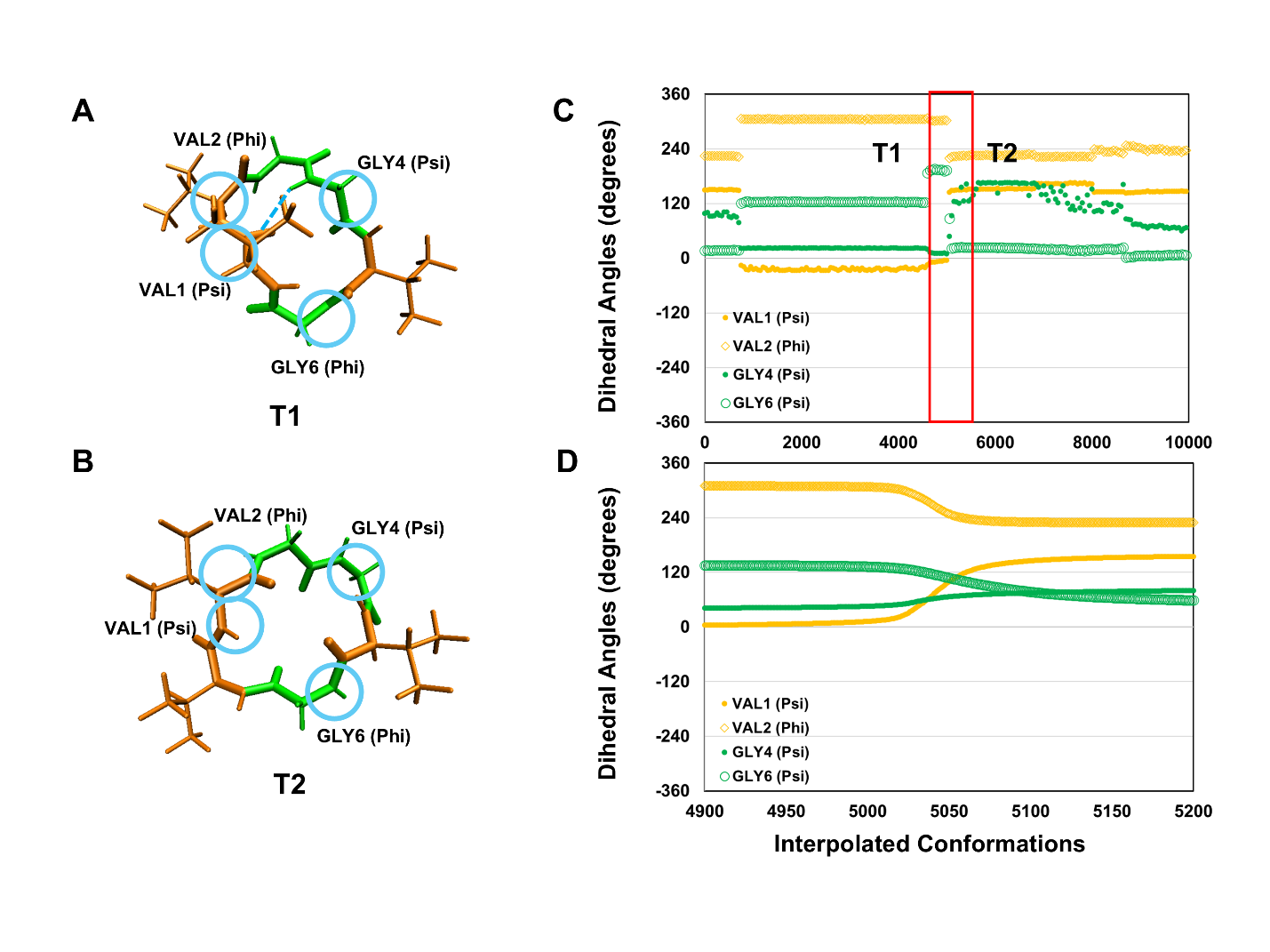

**Figure S3.** Closer observation of cylco-(VVGGVG) conformational transition from T1 to T2. **(A)** T1 of cyclo-(VVGGVG) from **Figure 5** **(B)** T2 of cyclo-(VVGGVG) from Figure 5 **(C)** Change in torsion angle over the interpolation pathway **(D)** Zoom in on the torsion angles during the transition between low energy state (T1) to high energy state (T2).

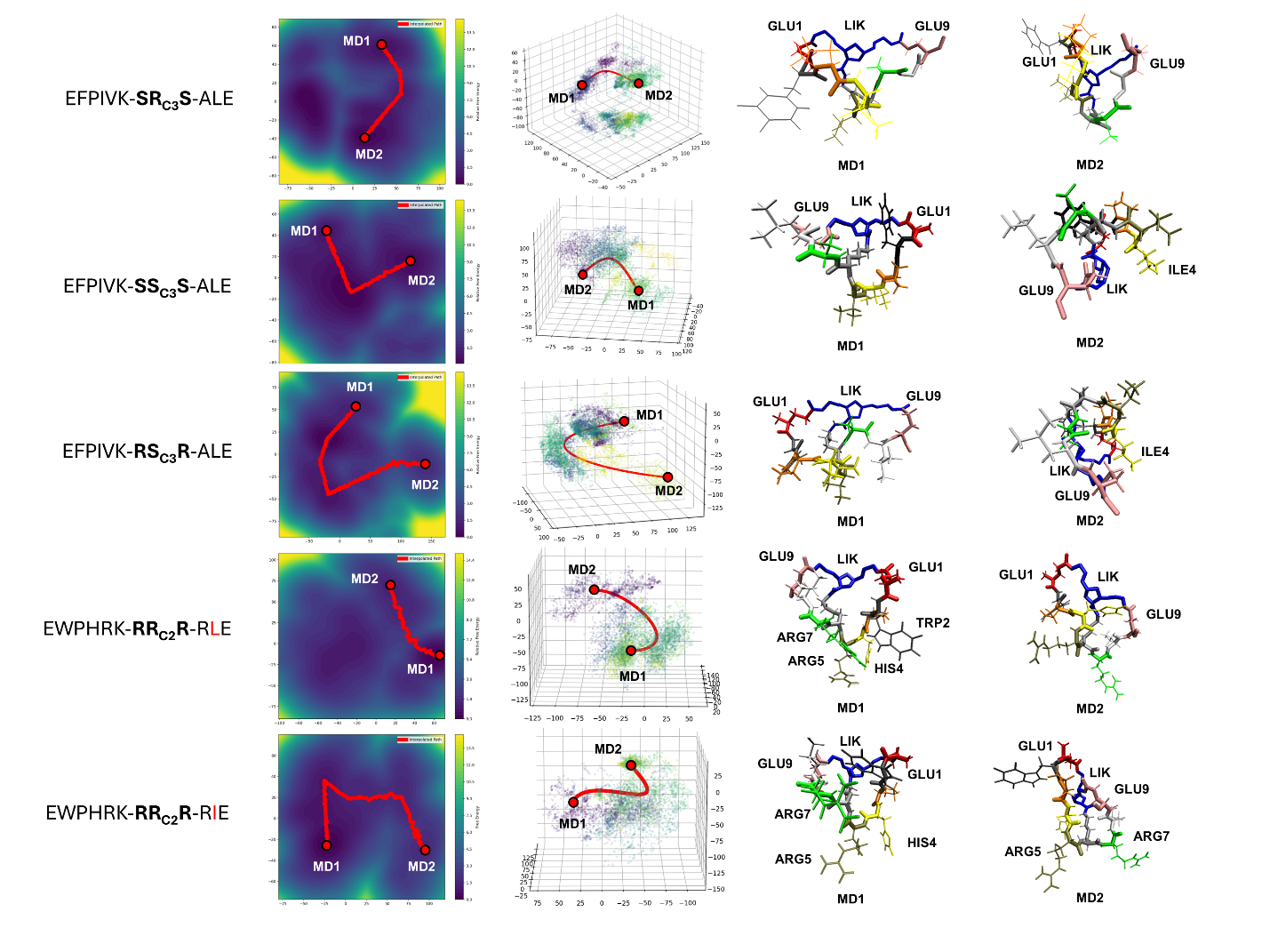

**Figure S4.** Minimum Energy Pathway (MEP) between two local energy minima, MD1 and MD2, of 2D probability contour map and 3D latent space of all five bicyclic peptides.

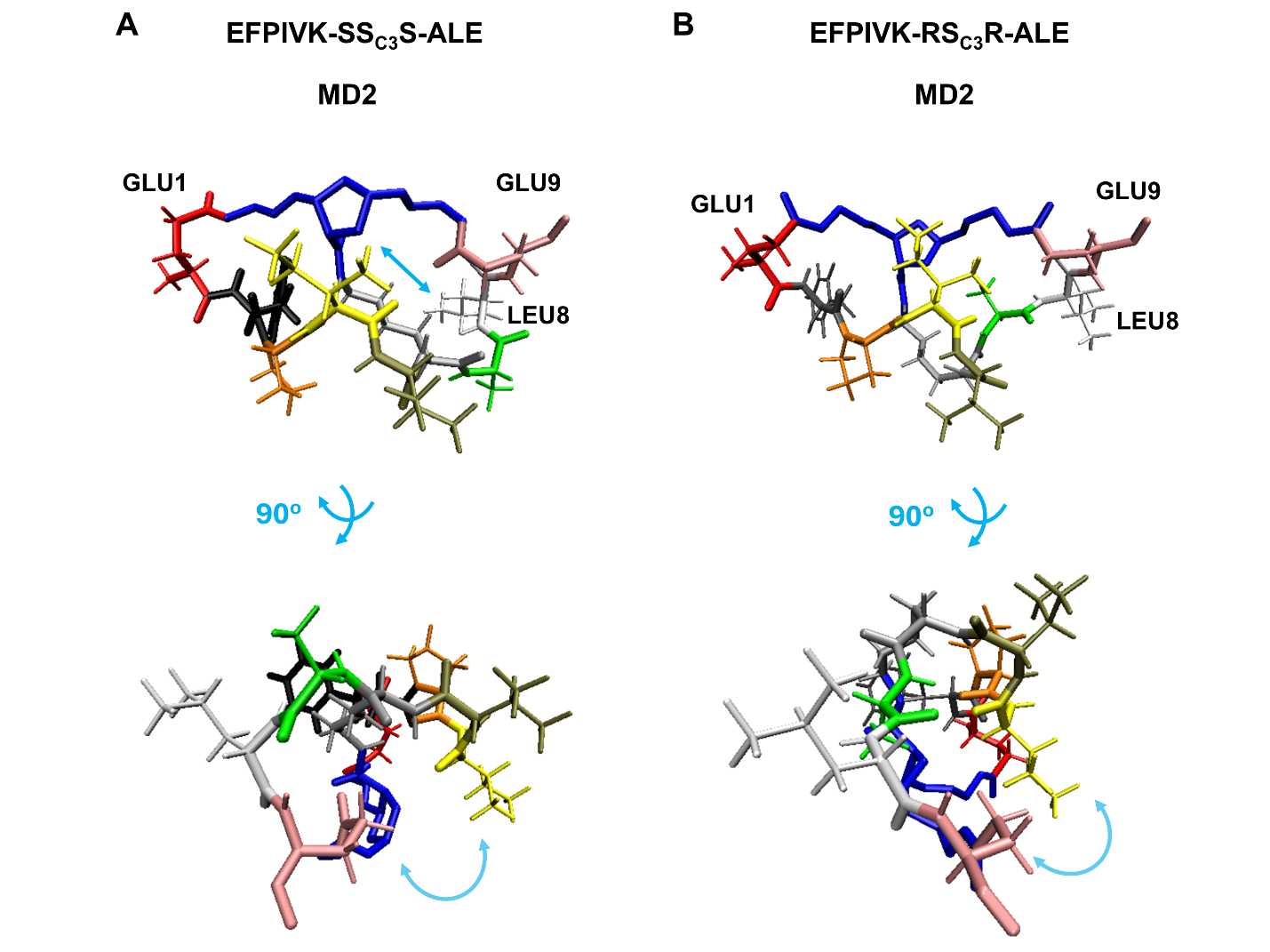

**Figure S5.** Different orientation of MD2 of **(A)** EFPIVK-SS_C3_S-ALE and **(B)** EFPIVK-RS_C3_R-ALE.

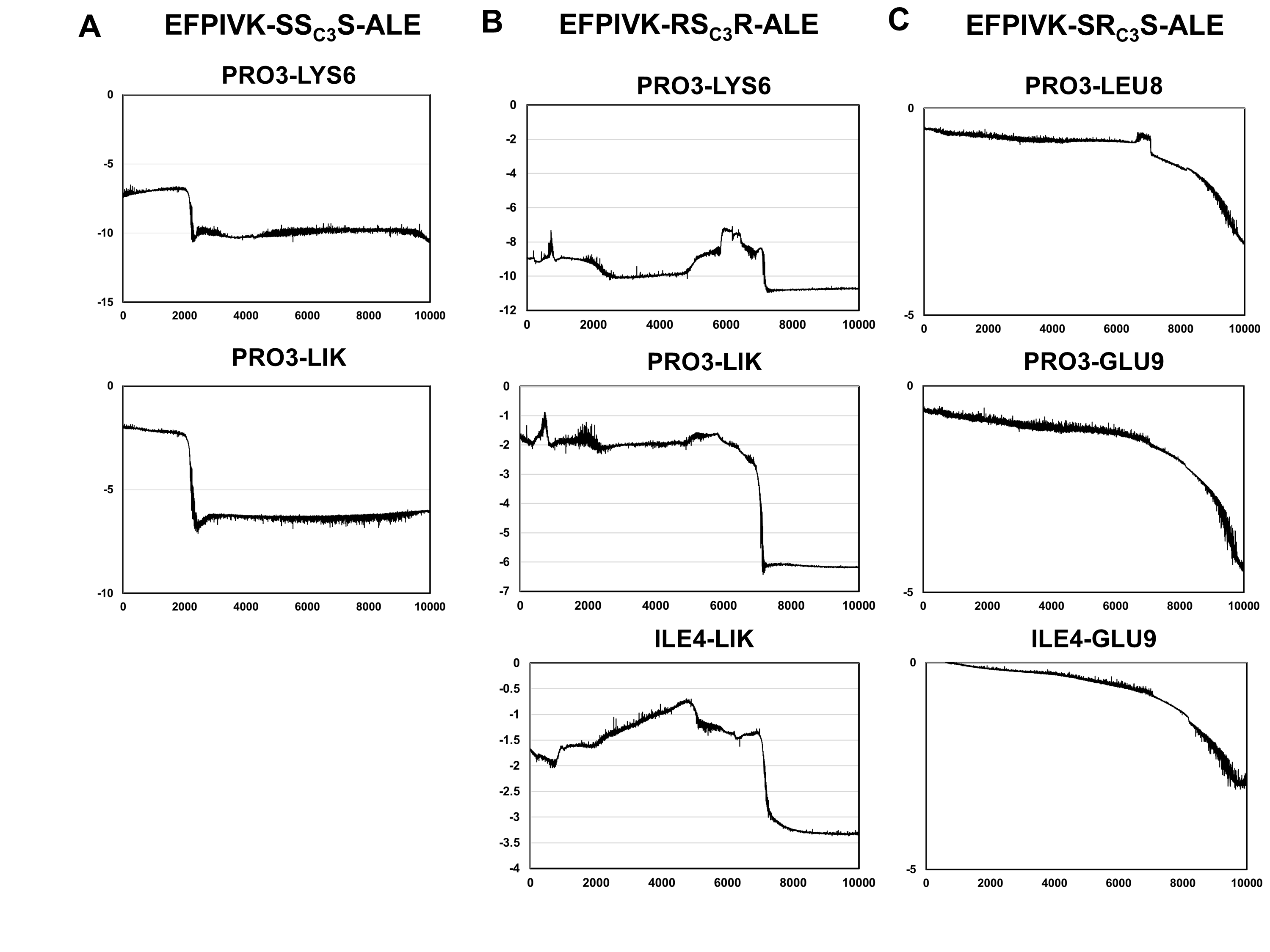

**Figure S6.** Strengthen interaction between PRO/ILE with surrounding residues during the conformational transition. (A) EFPIVK-SS_C3_S-ALE (B) EFPIVK-RS_C3_R-ALE (C) EFPIVK-SR_C3_S-ALE.

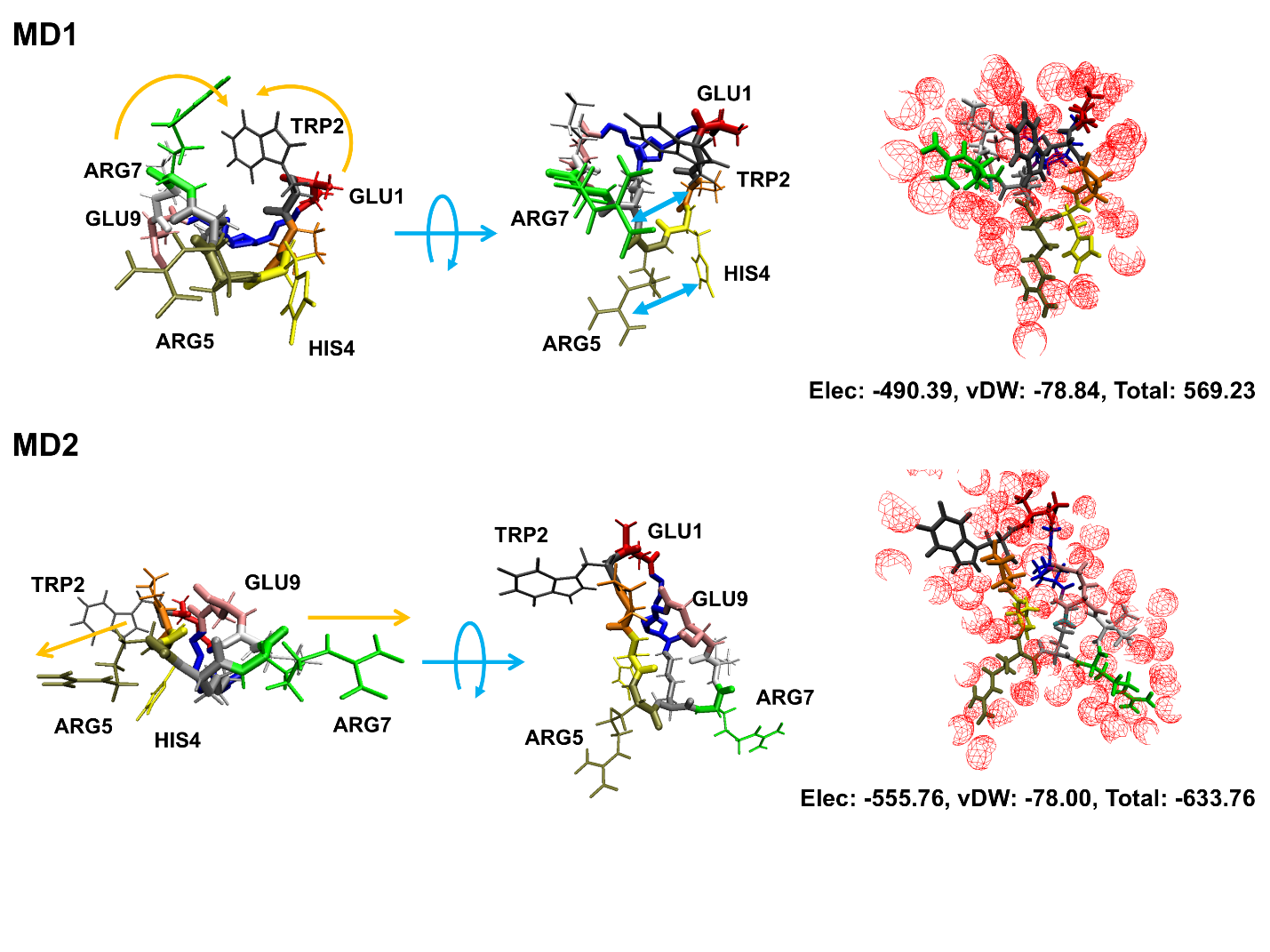

**Figure S7.** Protein-solvent interaction energy of distinct folded conformation of EWPHRK-RR_C2_R-RIE. MD1 with polar sidechain bury inward (orange curve) by forming cation-pi interaction (blue arrow) between ARG5 and HIS4 as well as ARG7 and TRP2 resulting in less favorable protein-solvent interaction. MD2 with polar sidechains is exposed (orange arrow) to the solvent resulting in more favorable protein-solvent interaction
